# X-PAIR: an ultrafast multitask framework for proteome-scale reconstruction of PPI networks and partner-specific interfaces from sequence

**DOI:** 10.64898/2026.07.20.739596

**Authors:** Sara Rescalli, Alessandra Carbone

## Abstract

Protein–protein interaction prediction and residue-level interface localisation are biologically intertwined but usually treated as separate computational problems. Here we present X-PAIR, a sequence-based multitask deep learning framework that jointly predicts whether two proteins interact and identifies their partner-specific interface residues. By combining protein language-model representations with lightweight cross-attention, X-PAIR requires neither structural templates nor multiple-sequence alignments. Across leakage-controlled benchmarks, it outperforms existing methods in both tasks, with substantial gains in interface localisation. Multitask learning preserves single-task performance while returning both outputs at near-single-task cost, enabling one million protein pairs to be analysed in under two hours—approximately 500-fold faster for interface prediction and 20-fold faster for PPI prediction than current approaches—thereby enabling proteome-scale analysis. Cross-species analyses reveal distinct evolutionary dependencies: interaction prediction benefits from multispecies training, whereas interface localisation remains robust across taxonomic scales. X-PAIR thus links proteome-scale interaction discovery to the residue-level determinants of partner-specific molecular recognition.

## INTRODUCTION

Protein–protein interactions (PPIs) lie at the core of virtually all cellular processes, orchestrating molecular pathways and assemblies that determine cellular function and behaviour. These interaction networks are fundamental not only for understanding basic biology but also for elucidating disease mechanisms, as perturbations in PPIs are implicated in a wide range of pathological conditions. Importantly, the disruption of PPIs is not a purely binary phenomenon: mutations can induce a spectrum of effects, ranging from subtle changes in binding affinity to a complete loss of interaction. In this context, residues located at interaction interfaces are particularly sensitive, as even minor alterations can propagate through the network and impair molecular recognition or complex stability. Critically, gaining mechanistic insight requires going beyond the binary question of whether two proteins interact, and instead addressing how and where binding occurs at the residue level, which is essential for rational drug design and therapeutic targeting. However, experimental characterization of PPIs remains challenging: existing techniques are costly, time-consuming, and incomplete even for well-studied organisms, with strong biases toward a limited set of model species. The situation is even more constrained for residue-level interface annotations, which typically depend on structural data that are far scarcer than sequence information. These limitations make computational approaches indispensable for large-scale inference of interactions and interfaces across diverse proteomes.

Protein partner discrimination and interface localisation are biologically related problems, since physical binding between two proteins is generally mediated by specific contact residues at their interface. Despite this biological coupling, most deep-learning approaches treat the two questions separately, focusing either on pair-level PPI prediction^1–3^ or on residue-level interface localisation^4–6^. In particular, when the goal is to characterize how two proteins interact, approaches ranging from docking-based pipelines^7^ to deep-learning interface predictors^8^ have achieved substantial progress, especially with recent advances in complex-structure prediction such as AlphaFold 3^9^, Boltz-2^10^, and RoseTTAFold2^11^. However, these approaches often require costly MSA construction and iterative complex-structure generation, making them computationally demanding and difficult to scale. Moreover, for a given candidate pair, complex-structure prediction models are primarily designed to generate a joint structural model rather than to explicitly predict a non-interaction class. Interaction status is therefore typically inferred post hoc from the predicted complex using scores that summarise inter-chain confidence and interface quality^12,13^. However, structural plausibility is not equivalent to binding specificity. While this issue was already recognized in the context of docking^14–16^, recent work shows that it persists in deep-learning complex predictors, which can assign plausible interfaces to non-cognate pairs^17–19^. Consistently, methods that explicitly optimise interface-aware signals for PPI discrimination can outperform post-hoc scores derived from predicted complexes^19,20^. Overall, however, the main limitation of these approaches remains the high computational cost of complex-structure prediction, making them better suited to rescoring selected candidates than to scalable *de novo* reconstruction of PPI networks across proteomes, where true interactions are intrinsically sparse.

On the other hand, some sequence-based binary PPI predictors have attempted to derive PPI contact maps as a secondary output. For example, D-SCRIPT^21^ and Topsy-Turvy^22^ extract contact-like maps from models trained primarily to distinguish interacting from non-interacting protein pairs. However, because these residue-level signals are not directly supervised as contacts, such models generally perform poorly as inter-protein contact predictors^23^. More recently, PEPpip^24^ applied post-hoc explainable AI to a sequence-based PPI predictor to generate more interpretable contact maps. Although it improves over D-SCRIPT and Topsy-Turvy, it remains less accurate than dedicated PPI-map predictors such as DRN-1D2D Inter^25^, which are trained explicitly with residue–residue contact supervision. Taken together, these observations indicate that, although partner discrimination and interface localisation are biologically coupled, neither task provides a fully sufficient proxy for the other: complex prediction can reveal plausible interfaces but does not, by itself, establish binding specificity, while binary PPI classifiers may capture residue-level signals without reliably resolving the full interaction interface. This motivates a more direct formulation in which PPI prediction is coupled with explicit residue-level interface supervision, while remaining lightweight enough to scale to large interactomes.

To bridge this gap, we introduce X-PAIR (Cross-attention Protein interAction and Interface pRediction), a lightweight and fast multitask architecture that jointly predicts whether two proteins interact and identifies the residues forming their binding interface. By leveraging pretrained protein language models (PLMs) and cross-attention mechanisms, X-PAIR captures both the global compatibility between protein pairs and local binding signals at the residue level, enabling interaction prediction and partner-specific interface localisation within a unified framework. Importantly, the model operates solely on amino acid sequences, without requiring structural templates or multiple sequence alignments, making it highly scalable and applicable to large-scale settings.

In this work, we make several key contributions. First, we introduce, to our knowledge, the first lightweight multitask architecture that jointly addresses both PPI prediction and residue-level interface identification within a single framework, without compromising performance compared to single-task models. Second, X-PAIR operates directly on paired amino acid sequences and requires neither structural information, structural templates nor multiple-sequence alignments, extending interaction and interface prediction across proteomes, including proteins lacking experimentally determined or reliably predicted structures. Third, X-PAIR produces partner-specific interface predictions, capturing context-dependent binding behaviour and addressing the challenge of signal disentanglement across multiple interaction partners. Fourth, its compact architecture, comprising relatively few trainable parameters, delivers substantial computational gains over leading task-specific approaches: X-PAIR is approximately 500-fold faster for interface prediction and 20-fold faster for PPI prediction, enabling one million protein pairs to be analysed in under two hours with modest computational resources. Fifth, we introduce curated datasets with compatible partitions across interaction and interface prediction tasks, enabling joint training and evaluation of both outputs. We release these resources to support the future development of multitask approaches. These datasets include a homology-controlled benchmark for assessing generalisation beyond closely related sequences and a dataset that enables systematic cross-species evaluation across evolutionary distances, revealing how evolutionary divergence affects predictive performance beyond sequence similarity alone.

## RESULTS

X-PAIR is a sequence-based multitask deep learning model designed to predict whether two proteins are interaction partners and identify the residues involved in their binding interface (Fig. 1). The model uses a frozen pretrained protein language model to encode the input sequences, followed by a shared trainable backbone that processes the paired residue-level representations and two task-specific prediction heads (Fig. 1A). The interaction prediction head outputs an interaction score representing the model’s confidence that the two proteins interact. When predictions are generated for a set of protein pairs, the resulting scores can be used to reconstruct a PPI network, where proteins are represented as nodes and predicted interactions as edges. Candidate interactions identified from the reconstructed network can then be further analysed by the interface prediction head, which assigns an interface score to each residue of both proteins, reflecting the model’s confidence that the residue contributes to the partner-specific interface. These residue-level predictions can be compared with known interface annotations when available and, optionally, projected onto an experimental or predicted structural model of the protein complex to support biological interpretation of the predicted interaction interface (Fig. 1B).

**Figure 1:**
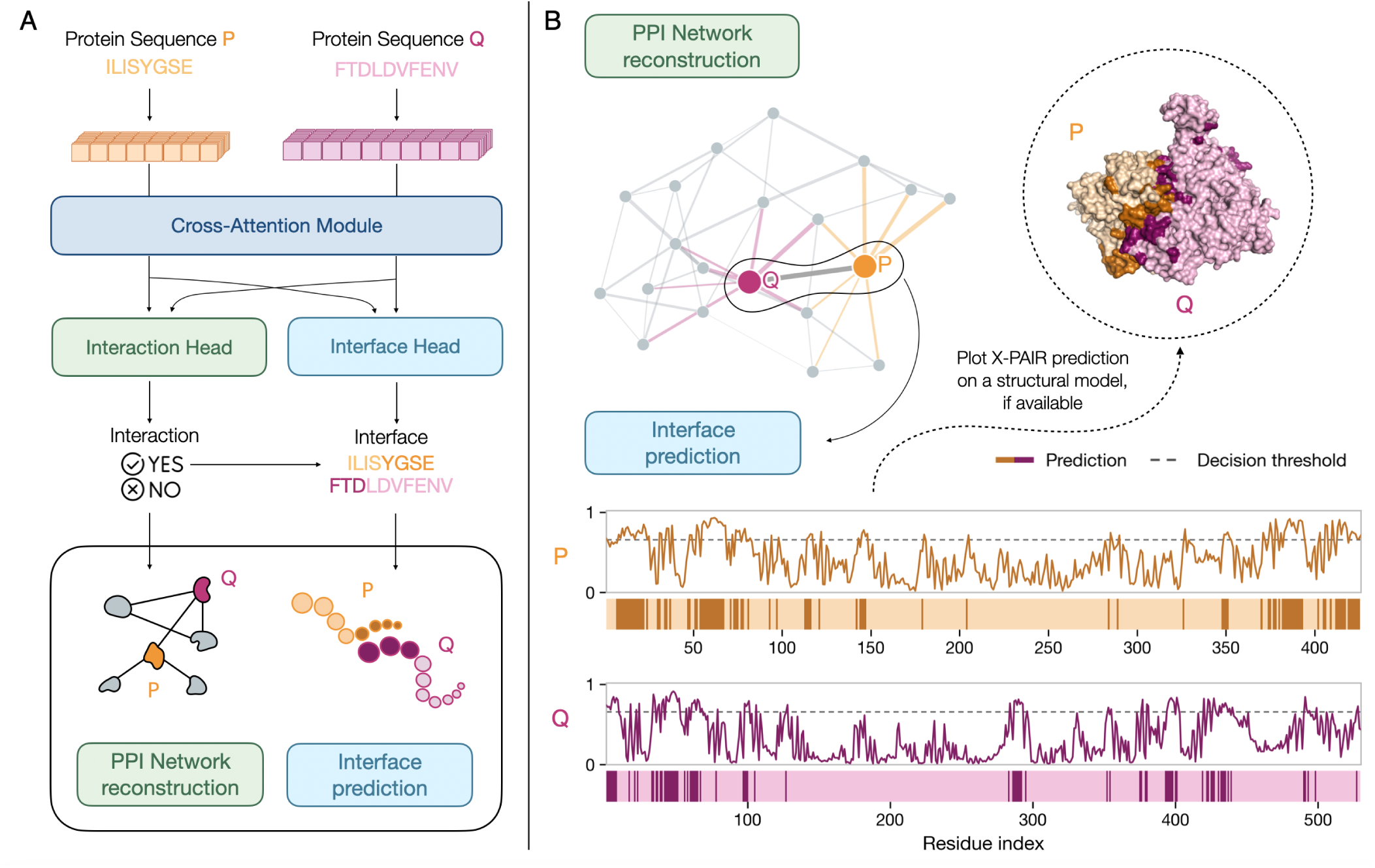
X-PAIR enables sequence-based prediction of PPIs and their binding interfaces. (A) Schematic overview of the X-PAIR architecture. Two protein sequences are first encoded into embeddings using a pretrained PLM and are then processed by a cross-attention module, which enables information exchange between the two candidate partners. Through two dedicated prediction heads, the model produces two complementary outputs: a pair-level interaction score and residue-level interface probabilities for both proteins. X-PAIR can therefore be used both for large-scale PPI network reconstruction and for residue-level inspection of selected protein interfaces. (B) Example application of X-PAIR to PPI network reconstruction and interface prediction. In the reconstructed network, nodes represent proteins and edges represent predicted interactions between protein pairs. Edge thickness is proportional to the interaction score assigned by X-PAIR, with thicker edges corresponding to higher-confidence interactions. For each predicted interacting pair, X-PAIR assigns an interface score between 0 and 1 to each residue, reflecting the model’s confidence that the residue contributes to the binding interface. A selected threshold can then be applied to these scores to identify predicted interface residues. When a structural model is available, these residue-level predictions can be mapped onto the three-dimensional protein structure to inspect the predicted interaction surface.

This design allows X-PAIR to be trained either in a multitask setting, where both prediction heads are optimised jointly, or in a single-task setting, where only the prediction head associated with the task under consideration is activated. Since no prior method simultaneously addresses protein partner identification and residue-level interface localisation within a unified framework, we first evaluated X-PAIR in single-task settings. Specifically, we benchmarked X-PAIR separately on well-established datasets for PPI prediction and residue-level interface prediction, enabling direct and fair comparison with existing task-specific methods. We then assessed X-PAIR in the joint multitask regime through complementary experiments designed to probe different aspects of generalisation. First, we tested whether the multitask model could generalise under a homology-controlled evaluation framework, limiting sequence identity between training and test protein pairs to 30% to reduce potential information leakage. Second, we performed cross-species analyses to investigate how predictive performance varies across organisms distributed along the evolutionary spectrum. In these experiments, we explicitly accounted for sequence similarity to the training set, enabling a controlled assessment of how evolutionary distance affects both protein partner identification and residue-level interface localisation.

### X-PAIR improves the performance for interface prediction

We first assessed X-PAIR in a single-task interface prediction setting by comparing its performance with several established sequence-based residue-level interface predictors. The evaluation included partner-specific methods such as BIPSPI+^5^, ECLAIR^6^, and PIONEER^4^, as well as non-partner-specific approaches including DLPred^26^, SCRIBER^27^, and DELPHI^28^. X-PAIR was benchmarked using the interface dataset introduced by Xiong et al. ^4^ , which provides partner-specific, residue-level interface annotations derived from experimentally determined co-crystal structures. Residues lacking interface annotations were excluded from both loss computation and evaluation, and protein pairs containing sequences longer than 2,000 amino acids were filtered out from the training and validation sets to ensure computational feasibility. During testing, all available protein pairs in the test set were evaluated regardless of sequence length.

As shown in Fig. 2A, X-PAIR achieves the best overall performance across most evaluation metrics. It obtains the highest AUROC (0.755) and AUPRC (0.353), with a particularly marked advantage in AUPRC over PIONEER, the second-best-performing method (AUPRC = 0.256). This improvement is particularly relevant in the strongly imbalanced setting of residue-level interface prediction, where non-interface residues predominate and AUPRC provides a more informative assessment of performance. X-PAIR also achieves the highest F1 score (0.397) and MCC (0.275), indicating a better balance between sensitivity and precision than the other sequence-based approaches. Its markedly higher recall (0.700) further reflects greater sensitivity in detecting interface residues and reduces the risk of missing biologically relevant binding sites. Although X-PAIR does not achieve the highest accuracy, this metric is threshold-dependent and less informative under class imbalance. Remarkably, despite relying exclusively on sequence information, X-PAIR outperforms the structure-informed versions of both ECLAIR and PIONEER across all metrics except AUROC and accuracy.

**Figure 2:**
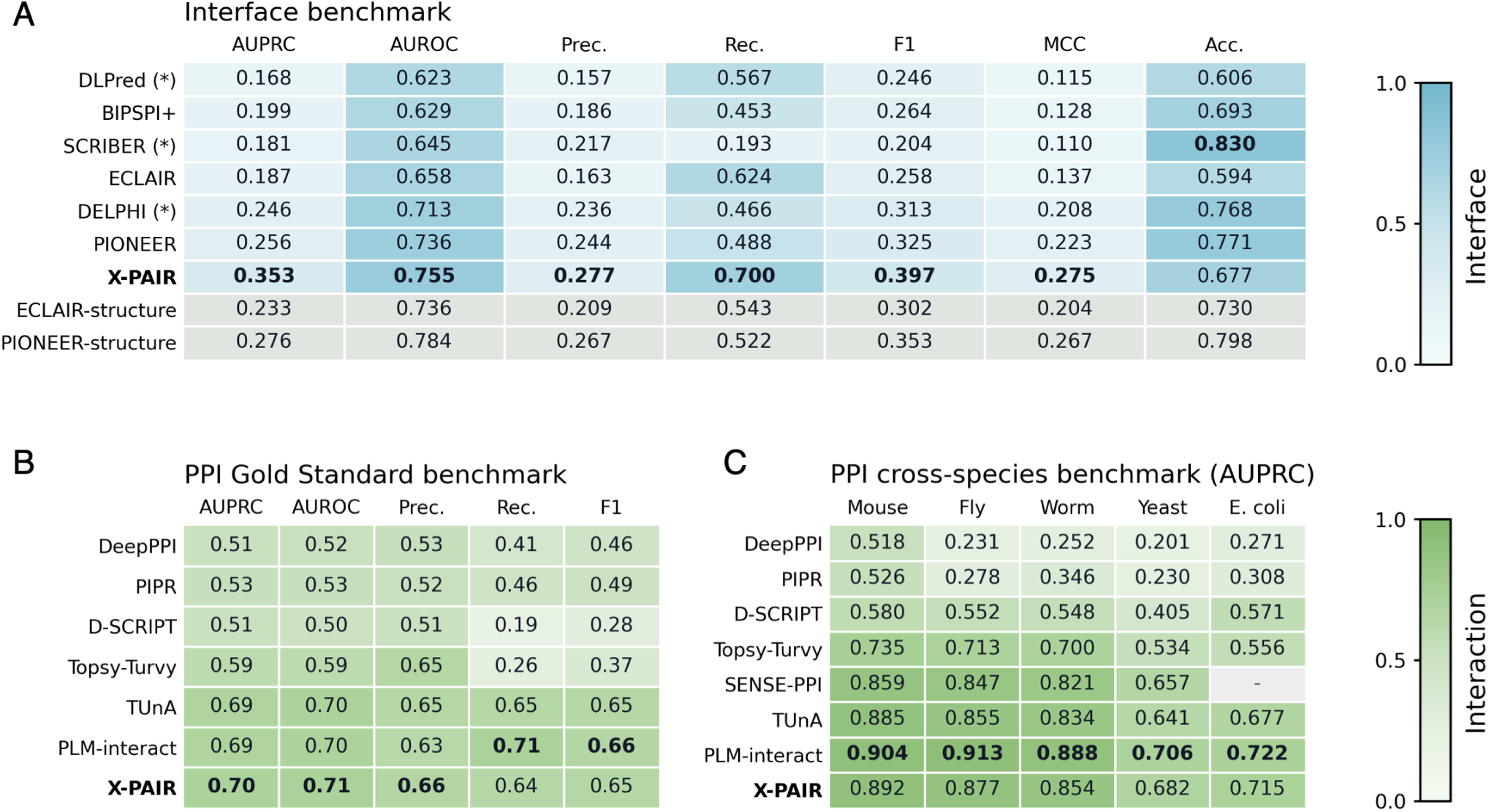
Benchmarking single-task X-PAIR models against existing sequence-based interface and PPI prediction methods. (A) Comparison of interface prediction performance on the benchmark dataset introduced by Xiong et al. ^4^ . Baseline metrics were taken from Supplementary Tables 2 and 3 of the PIONEER study^4^, whereas X-PAIR results were obtained in this work. Methods marked with an asterisk (*) are single-chain predictors, whereas BIPSPI+, ECLAIR, PIONEER, and X-PAIR are partner-specific methods. For reference, the structure-informed versions of ECLAIR and PIONEER are also included in the comparison. (B) Comparison of PPI prediction performance on the Gold Standard benchmark dataset^29^. Baseline results were taken from the PLM-interact study^1^, whereas X-PAIR results were obtained in this work. (C) Cross-species PPI prediction performance on the benchmark introduced by Sledzieski et al. ^21^ . Models were trained on human protein pairs and evaluated on non-human species. Performance is reported as AUPRC. SENSE-PPI values were taken from the SENSE-PPI study^3^, whereas all other baseline values were taken from the PLM-interact study^1^. X-PAIR results were obtained in this work.

X-PAIR is also substantially more computationally efficient than PIONEER. Specifically, X-PAIR processes approximately 150 protein pairs per second, whereas PIONEER has a reported inference time of more than 3 seconds per protein pair^4^.

### X-PAIR reaches state-of-the-art performance for PPI prediction

We evaluated X-PAIR in the single-task interaction prediction setting by comparing its performance with several established sequence-based PPI prediction methods, including PIPR^30^, DeepPPI^31^, D-SCRIPT^21^, Topsy-Turvy^22^, SENSE-PPI^3^, TUnA^2^, and PLM-interact^1^.

Evaluation was first conducted on the leakage-free human Gold Standard dataset curated by Bernett et al. ^29^ . This benchmark is explicitly designed to minimise sequence similarity across training, validation, and test splits, thereby reducing the risk of information leakage due to homologous proteins. For computational reasons, protein pairs containing sequences longer than 2,000 amino acids were excluded from the training and validation splits, while evaluation was conducted on the full test benchmark. Fig. 2B reports the comparison with existing interaction prediction methods. X-PAIR achieves the highest AUPRC (0.70) and AUROC (0.71), indicating strong discriminative ability. In terms of precision and recall, X-PAIR maintains a well-balanced trade-off (0.66 precision, 0.64 recall). Overall, these results demonstrate competitive or superior performance across all major metrics, confirming the effectiveness of the proposed framework in the single-task interaction setting.

We then extended the analysis to cross-species generalisation using the cross-species benchmark introduced by Sledzieski et al. ^21^ , which is specifically designed to assess transfer performance in PPI prediction. Under this protocol, models were trained exclusively on human PPI data and subsequently evaluated on five evolutionarily distinct species: *Mus musculus*, *Drosophila melanogaster*, *Caenorhabditis elegans*, *Saccharomyces cerevisiae*, and *Escherichia coli*. This setup enables a systematic investigation of how predictive performance degrades with increasing evolutionary distance from the training distribution. X-PAIR outperforms all evaluated baseline models except PLM-interact, with which it achieves comparable performance across all considered species (Fig. 2C). In general, compared to earlier approaches such as DeepPPI, more recent architectures show reduced sensitivity to phylogenetic distance from the human training distribution.

Importantly, although PLM-interact achieves performance comparable to X-PAIR, it is substantially less computationally efficient. Processing one million protein pairs requires approximately two days with PLM-interact, compared with about two hours with X-PAIR, making X-PAIR roughly 20 times faster and considerably more suitable for large-scale applications.

### The multitask framework of X-PAIR efficiently predicts both interactions and interfaces

To rigorously evaluate the multitask performance of X-PAIR, we introduced the X-fair dataset, specifically designed to reduce information leakage between training, validation, and test sets. This is particularly important in biological prediction tasks, where the presence of highly similar proteins across data splits can lead to overestimated generalisation performance^29^. Our X-fair dataset includes interface data from multiple species and interaction data from four species: *Homo sapiens*, *Gallus gallus*, *Saccharomyces cerevisiae*, and *Drosophila melanogaster*. All proteins were first clustered at 30% sequence identity, and each cluster pair was then assigned in its entirety to the training, validation, or test set (see Methods), thereby reducing homology-based leakage and enabling a stringent evaluation of generalisation across both tasks.

Fig. 3A shows the performance of X-PAIR on the X-fair dataset, comparing the multitask model with its single-task counterparts trained separately for interaction prediction and interface localisation. All models were trained using the same task-specific data splits and hyperparameters, allowing a fair assessment of the effect of joint optimization. Exact metric values are reported in Table S5. Overall, X-PAIR achieves strong performance on both interaction prediction and residue-level interface prediction. As expected, performance is higher for the interaction prediction task (the distributions of predicted interaction probabilities for positive and negative test pairs are shown in Fig. S2), whereas interface prediction remains more challenging. This is likely due both to the smaller amount of PDB-derived data available for training interface annotations and to the intrinsic difficulty of the task, which requires residue-level predictions and is therefore highly sensitive to small changes in the binding of the protein. Importantly, performance in the multitask setting remains comparable to that obtained with single-task training. This shows that jointly optimizing the two objectives does not lead to a degradation in predictive quality, suggesting that the multitask model can exploit shared sequence representations across the two levels of prediction without sacrificing accuracy on either task. The agreement between the single-task and multitask models is further illustrated in Fig. S3.

**Figure 3:**
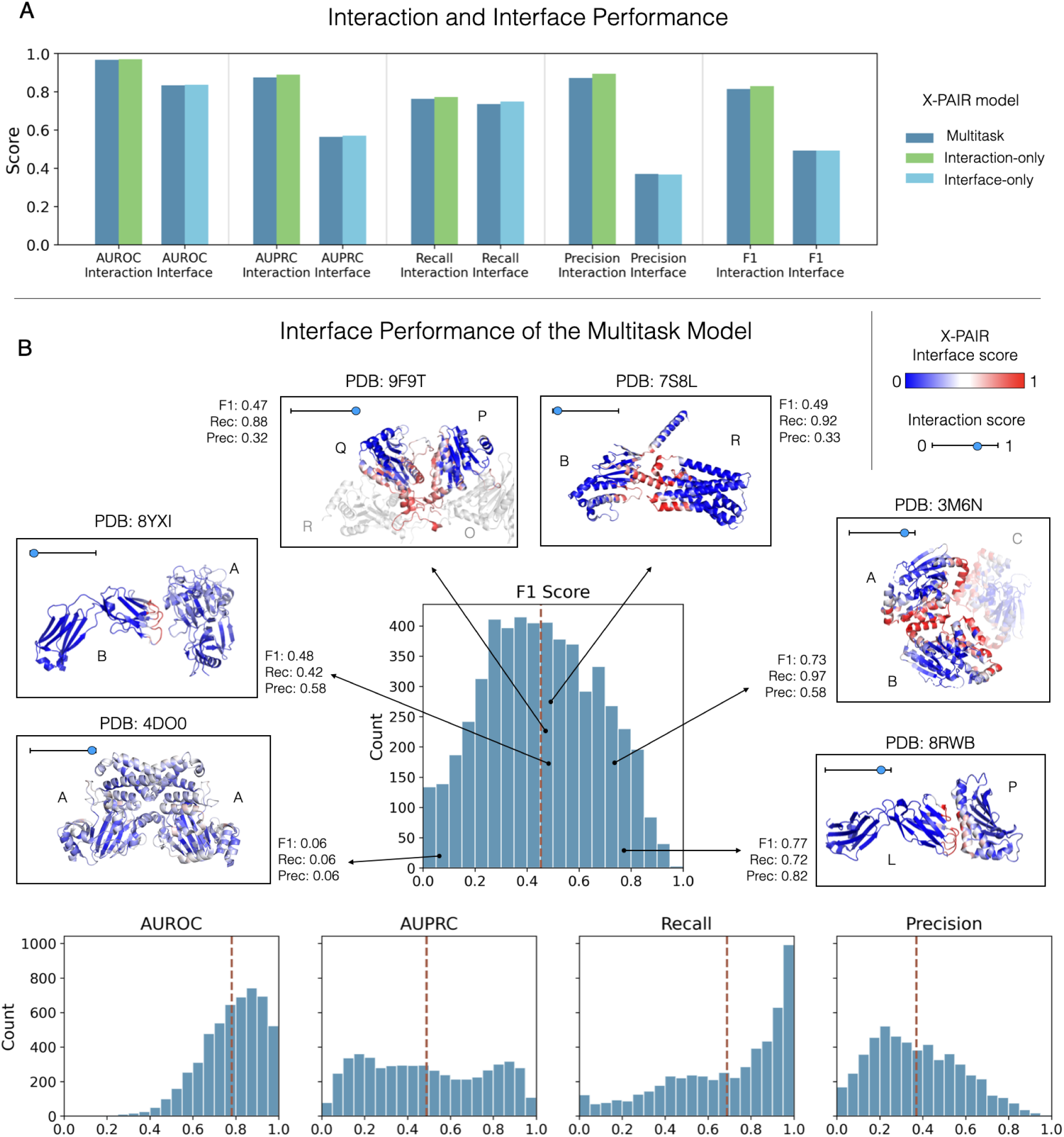
Evaluation of X-PAIR multitask model. (A) Comparison of the multitask model with the corresponding single-task baselines, namely the interaction-only and interface-only models. Performance is reported separately for the interaction prediction task and for the interface prediction task, showing that the multitask setting achieves competitive results on both objectives. The decision threshold for precision, recall, and F1 score is always set to 0.5. AUPRC is computed as average precision. (B) Interface prediction performance of the multitask model. Distribution of pair-level interface prediction performance across test pairs; red dashed lines indicate the mean value for each metric. Representative examples are also shown by mapping X-PAIR interface predictions onto the corresponding experimental PDB structures, allowing a qualitative comparison between predicted and true interfaces. For each pair, the corresponding F1 score, precision, and recall are reported. The interaction score predicted by X-PAIR is also reported for each pair in the left corner of each box. Structural figures were generated with PyMOL^32^.

We therefore focused specifically on the performance of the X-PAIR multitask model in interface prediction, with Fig. 3B showing the distributions of the main evaluation metrics across protein pairs in the test set. Overall, AUROC values are high for most pairs, indicating that the model generally ranks interface residues above non-interface residues. However, because interface residues represent only about 17% of the dataset, AUROC alone does not fully capture performance under class imbalance, making complementary metrics such as AUPRC, precision, and recall more informative. As expected, AUPRC values are more broadly distributed, reflecting differences in difficulty across protein pairs. Nevertheless, their mean remains clearly above the random baseline of approximately 0.17, indicating that the model captures meaningful residue-level signals despite the strong class imbalance. Although precision is more variable, recall remains consistently high. This profile is well suited to interface identification, where missing true interface residues may be more problematic than predicting a broader set of candidate sites, particularly when predictions are used to guide downstream experiments. Moreover, some residues classified as false positives may correspond to genuine interaction sites that have not yet been experimentally characterized, potentially leading to an underestimation of precision.

### Residue-level interface predictions reveal interpretable binding patterns beyond pair-level confidence

We next examined a small set of representative protein pairs to qualitatively explore how pair-level interaction scores and residue-level interface predictions behave across different cases, and how the spatial distribution of predicted interfaces can provide information beyond evaluation metrics. To this end, we examined six representative protein pairs for which X-PAIR interface predictions (Fig. S4) were mapped onto the corresponding experimental PDB structures (Fig. 3B). These examples were selected from X-fair interface test set to span a broad range of residue-level performance, from strong interface localisation to intermediate and difficult cases, and were analysed together with the interaction prediction scores assigned by the model.

These examples show that residue-level interface prediction and pair-level interaction probability do not follow a simple one-to-one relationship. Four of the six protein pairs (PDB 8RWB, PDB 3M6N, PDB 9F9T, and PDB 4DO0) received high interaction scores, indicating that X-PAIR recognized them as likely interaction partners even when residue-level localisation was imperfect. Conversely, lower interaction scores were observed for two of the complexes (PDB 8YXI and PDB 7S8L), despite the fact that their residue-level predictions retained interpretable structural features. Notably, both represent atypical interaction contexts that may weaken the global sequence signal required for interaction recognition: PDB 8YXI is a cross-species mouse–virus complex, whereas chain B in PDB 7S8L is an engineered chimeric G protein rather than a naturally occurring partner. Together, these observations suggest that interaction prediction and interface localisation rely, at least in part, on different types of information and should therefore be interpreted as related but non-equivalent components of the overall prediction task, as further developed in the Discussion.

We next examined these examples at residue-level resolution to determine whether the spatial distribution of predicted interface residues provides biologically interpretable information beyond aggregate performance scores. In the best-performing cases, X-PAIR produces compact and well-focused predictions over structurally meaningful binding surfaces. For example, in PDB 8RWB, which comprises a human immune ligand bound to the light chain of a blocking antibody^33^, the model reaches an F1 score of 0.77. Similarly, for the self-interaction of a bacterial quorum-sensing enzyme in PDB 3M6N^34^, X-PAIR achieves an F1 score of 0.73. Because this enzyme forms a homotrimer, mapping the predictions onto a single chain pair reveals additional predicted interface regions that actually coincide with the position occupied by the third subunit. This is consistent with our definition of the interface, which combines all observed interaction configurations for a given protein pair, including contacts involving repeated or symmetry-related subunits.

Examples with intermediate F1 scores of approximately 0.48 further show that similar aggregate performance can arise from distinct but interpretable prediction patterns. In both PDB 7S8L and PDB 9F9T, X-PAIR achieves relatively high recall but lower precision, although for different reasons. PDB 7S8L comprises a peptide-activated human membrane receptor bound to an engineered G protein that stabilises its active signaling state^35^. Most true interface residues are recovered, whereas the additional predicted residues remain spatially close to the annotated interface in the three-dimensional structure, broadening the predicted binding region rather than forming unrelated surface patches. In PDB 9F9T, which contains neighbouring proteasome subunits from *Trypanosoma cruzi* ^36^, the partner-specific interface is also recovered, but some additional predictions occur in regions that contact adjacent subunits within the multimeric assembly. PDB 8YXI, involving a viral surface protein bound to the light chain of a mouse neutralizing antibody^37^, shows a different pattern. Precision and recall are more balanced overall, but performance differs between the two partners, with better recovery of the antibody-side interface than of the corresponding viral surface patch. Thus, intermediate F1 scores do not necessarily indicate uniformly poor predictions but may reflect spatially coherent and biologically meaningful interaction patterns.

Other cases remain substantially more challenging. PDB 4DO0 represents the homomeric assembly of a human histone demethylase in a structure determined in the presence of the small-molecule ligand daminozide. Although X-PAIR correctly identifies the buried protein core as non-interacting, it fails to localise the specific surface region corresponding to the annotated interface, resulting in an F1 score of 0.06. This case therefore represents a genuine failure of residue-level interface localisation and highlights a limitation of the model.

### Interaction and interface prediction show distinct cross-species generalisation patterns

To further investigate the multitask capabilities of X-PAIR, we evaluated its performance from an evolutionary perspective, assessing its ability to generalise to phylogenetically distant species. To this end, we constructed the X-human dataset, in which training data were restricted to *Homo sapiens* proteins for both interaction and interface prediction tasks. This setting enables a controlled analysis of how evolutionary distance affects model performance across different test species, including *Escherichia coli*, *Arabidopsis thaliana*, *Caenorhabditis elegans*, *Bombyx mori*, *Danio rerio*, *Oryctolagus cuniculus*, *Sus scrofa*, and *Bos taurus*.

Fig. 4A shows how the model’s MCC varies across species as a function of their average sequence identity to the training set; the corresponding AUROC and AUPRC results are reported in Fig. S5. Pair sequence identity to the training set was quantified by comparing each test protein against all training proteins using MMseqs2^38^, retaining the maximum identity value for each test protein, and then averaging these values across all test proteins from the same species (see Methods). For interaction prediction, we observe a clear decrease in performance as similarity to the training set decreases. This trend suggests that the interaction model relies, at least in part, on sequence-level proximity between the test proteins and the training distribution, with less similar species representing a more challenging generalisation setting. In contrast, for interface prediction, no clear decline in MCC is observed as sequence similarity decreases. This represents an interesting difference between the two tasks and may suggest that interface prediction is less directly driven by sequence similarity. Moreover, interface prediction exhibits overall lower performance than interaction prediction, reflecting both the intrinsic difficulty of the task and the substantially smaller amount of training data available.

**Figure 4:**
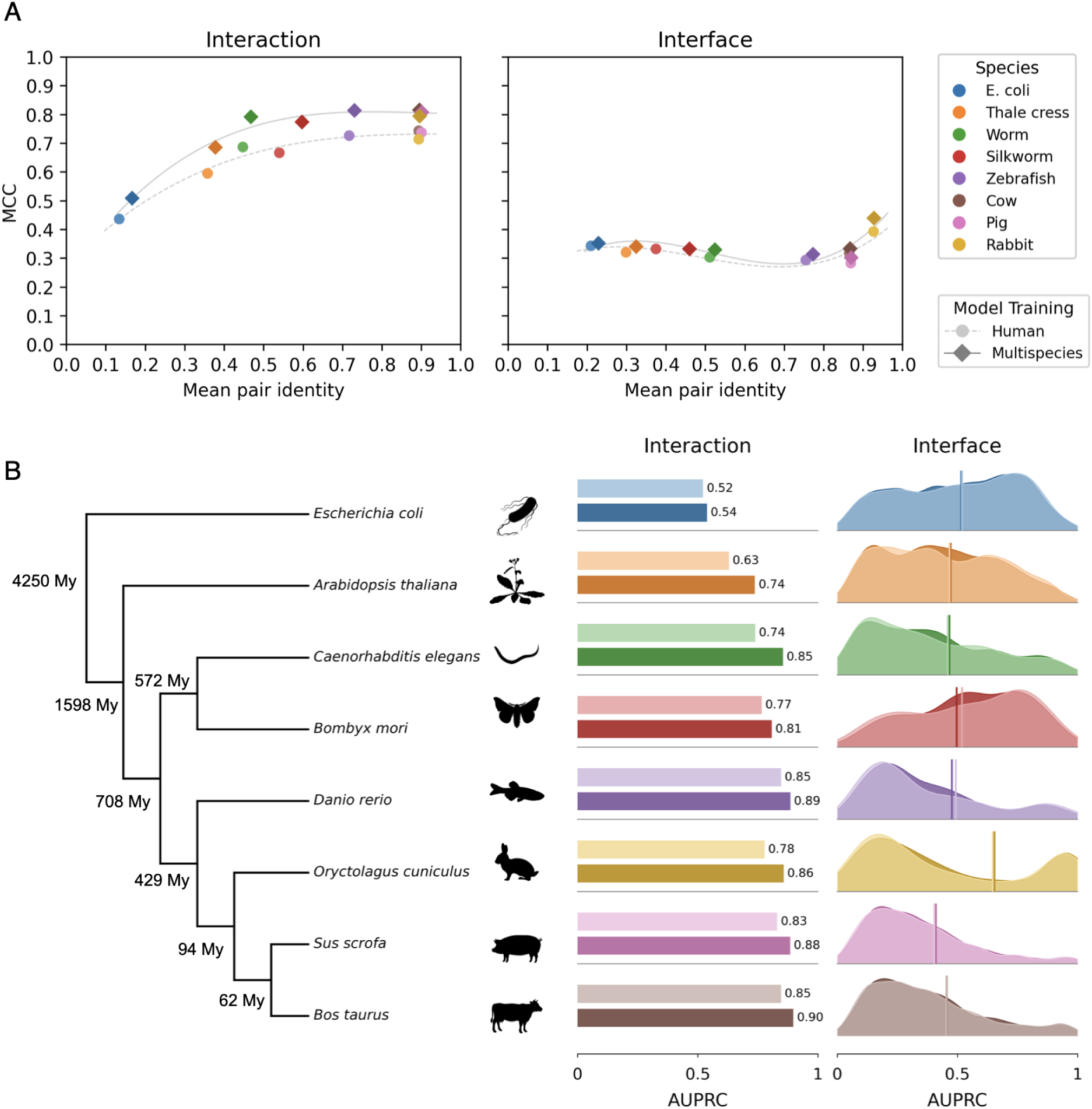
Cross-species generalisation of interaction and interface prediction. (A) Matthews correlation coefficient (MCC) as a function of mean pairwise sequence identity between test proteins and the training set, evaluated across multiple test species. Results are reported for both interaction prediction at the protein-pair level and interface prediction at the residue level, comparing a model trained exclusively on human data (X-human dataset) with a model trained on multiple species (X-multispecies dataset) comprising human, chicken, fly, and yeast proteins. (B) Area under the precision–recall curve (AUPRC) for the same two models across test species, shown as a function of evolutionary distance, measured in millions of years. For interface prediction, the distribution of AUPRC values across protein pairs is shown. Species icons were obtained from PhyloPic (https://www.phylopic.org).

Motivated by previous observations in SENSE-PPI^3^, where adding species evolutionarily closer to the target test species improved interaction prediction performance, we constructed an additional dataset, named X-multispecies. In this setting, selected non-human species were included in the training set together with human proteins. Specifically, training was performed on proteins from *H. sapiens*, *G. gallus*, *D. melanogaster*, and *S. cerevisiae*. For interaction prediction, multispecies training consistently improves performance across all test species in terms of MCC, as shown in Fig. 4A. As expected, for each species this improvement is accompanied by a small increase in sequence similarity to the training set. However, this increase in similarity is modest compared with the observed gain in predictive performance. This suggests that the improvement obtained with multispecies training cannot be explained solely by the presence of more similar sequences in the training data, but may instead reflect the ability of the model to exploit broader cross-species information. For interface prediction, instead, this pattern is not observed. Moving from human-only training to multispecies training does not lead to a substantial improvement in MCC. This reinforces the idea that interface prediction may follow a different generalisation regime from interaction prediction.

We then asked whether a similar behaviour could be observed when performance is analysed across different levels of evolutionary divergence rather than as a function of sequence similarity alone. Fig. 4B shows the performance of the same models across species ordered along a broad phylogenetic scale. Overall, interaction prediction shows a tendency toward lower performance as evolutionary divergence increases. This suggests that the interaction model may be affected by the evolutionary context of the test species, with more divergent branches of the tree of life generally representing a more challenging generalisation setting. The behaviour of interface prediction is again different. No clear trend is observed along this evolutionary scale. Indeed, some highly divergent species, such as *E. coli*, show better interface prediction performance than vertebrate species such as *S. scrofa*. This suggests that evolutionary divergence alone is not sufficient to explain interface prediction performance, and that other factors may play a stronger role in this task.

Taken together, these results indicate that interaction prediction and interface prediction behave differently under cross-species generalisation. Interaction prediction is strongly affected by sequence similarity, evolutionary distance, and the taxonomic composition of the training set, and it benefits substantially from multispecies training. By contrast, interface prediction shows lower overall performance, no clear dependence on sequence similarity or evolutionary distance, and little improvement when non-human species are added to the training set. Although the two tasks are biologically connected, these observations suggest that a black-box model may exploit different types of information for each task. In particular, interaction prediction appears to rely more directly on global sequence-level signals, whereas interface prediction may depend on more local determinants of the interacting regions, such as the conservation of interface residues, local structural or physicochemical compatibility, and the availability and quality of residue-level annotations.

### Interface prediction remains stable across evolutionary distances and taxonomic groups

Motivated by the observation that interface prediction performance remains stable even at low sequence similarity to the training set, we further investigated the ability of X-PAIR to generalise across species at different evolutionary scales. More precisely, we analysed interface prediction performance as a function of both evolutionary divergence and sequence identity to the training data. To this end, we constructed the X-taxonomic-interface dataset, as described in the Methods, and evaluated the single-task version of X-PAIR exclusively on the interface prediction task. For each taxonomic group of interest, the training set was restricted to interaction pairs in which neither partner belongs to the target group, while the test set included all pairs with at least one protein from that group. We first used *Mammalia* as a case study, training on interaction pairs involving only non-mammalian proteins and evaluating performance exclusively on mammalian proteins in the test set, regardless of the species of their interaction partners. Non-mammalian partners in the test set may therefore have been observed during training, and in fact they were excluded from the evaluation. In addition, a mammalian protein under evaluation may have been indirectly observed during training if an identical sequence was present in a non-mammalian species.

Fig. 5A shows, across different mammalian species, how mean AUROC performance varies as a function of evolutionary divergence time and sequence identity to the training set. For each species, evolutionary distance was estimated using the TimeTree resource^39^, by considering the divergence time from its closest sister lineage (see Methods). Sequence identity to the training data was instead computed using MMseqs2 (see Methods), by retaining, for each test protein, the maximum sequence identity observed against all proteins in the training set, and then averaging these values at the species level. A regression line with a 95% confidence interval was added to the plot to model the relationship between mean AUROC and evolutionary distance from the present, providing a species-level estimate of this trend. The same analysis was subsequently extended to other major lineages, including *Viridiplantae* (Fig. 5B), *Viruses* (Fig. 5C–Fig. 5D), *Archaea*, *Fungi*, and *Insecta* (Fig. S6).

**Figure 5:**
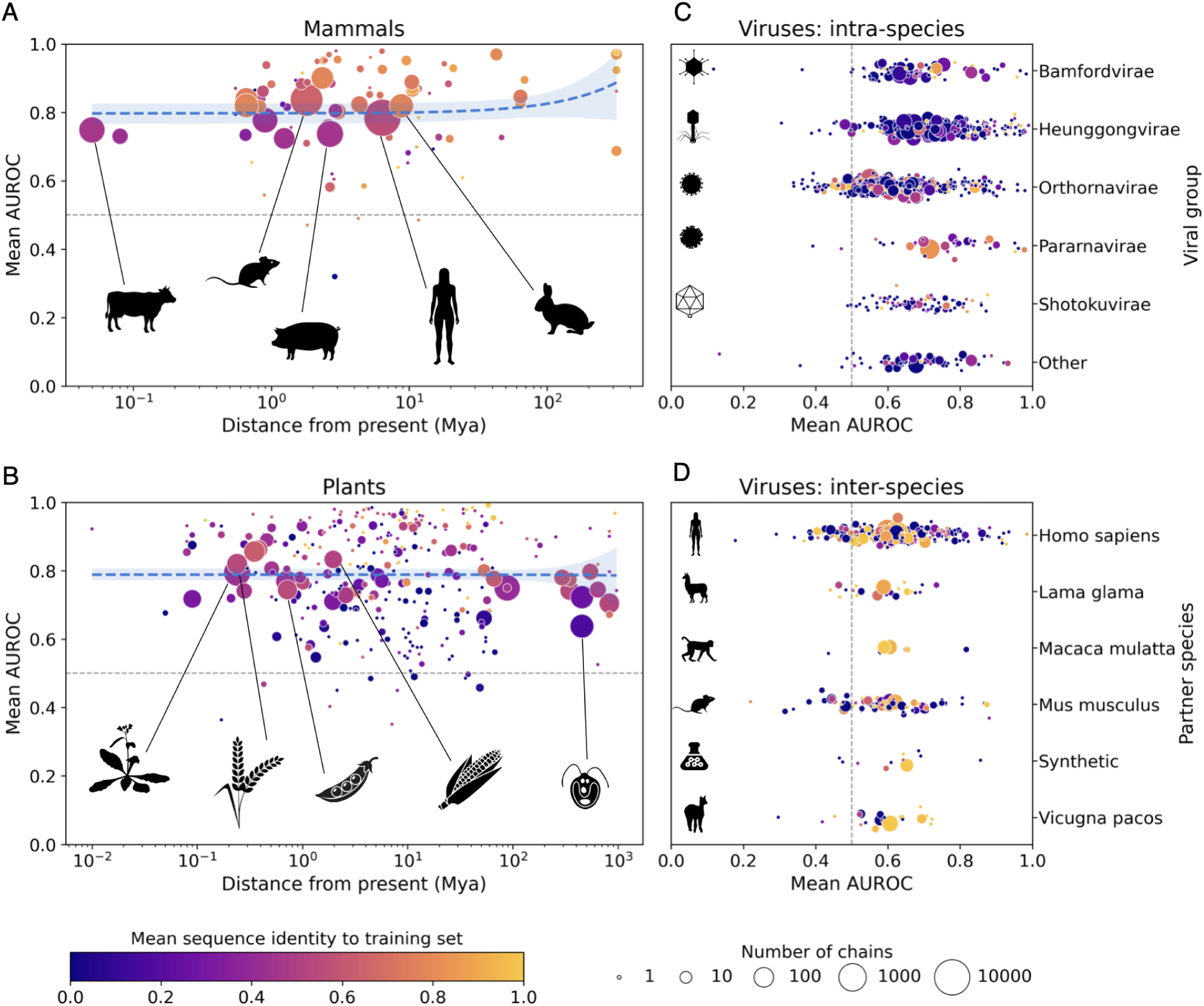
Performance across species as a function of evolutionary divergence time and sequence identity to the training set. Each point represents a species; marker size is proportional to the number of PDB chains, and colour denotes the mean sequence identity to the training set. (A) Mean AUROC as a function of evolutionary divergence time, measured in millions of years, for *Mammalia*. (B) Same as in (A) for *Viridiplantae*. (C) Mean AUROC for viral proteins involved in intra-species interactions, grouped by viral group. (D) As in (C) for inter-species interactions, grouped by partner species.

**Figure 6:**
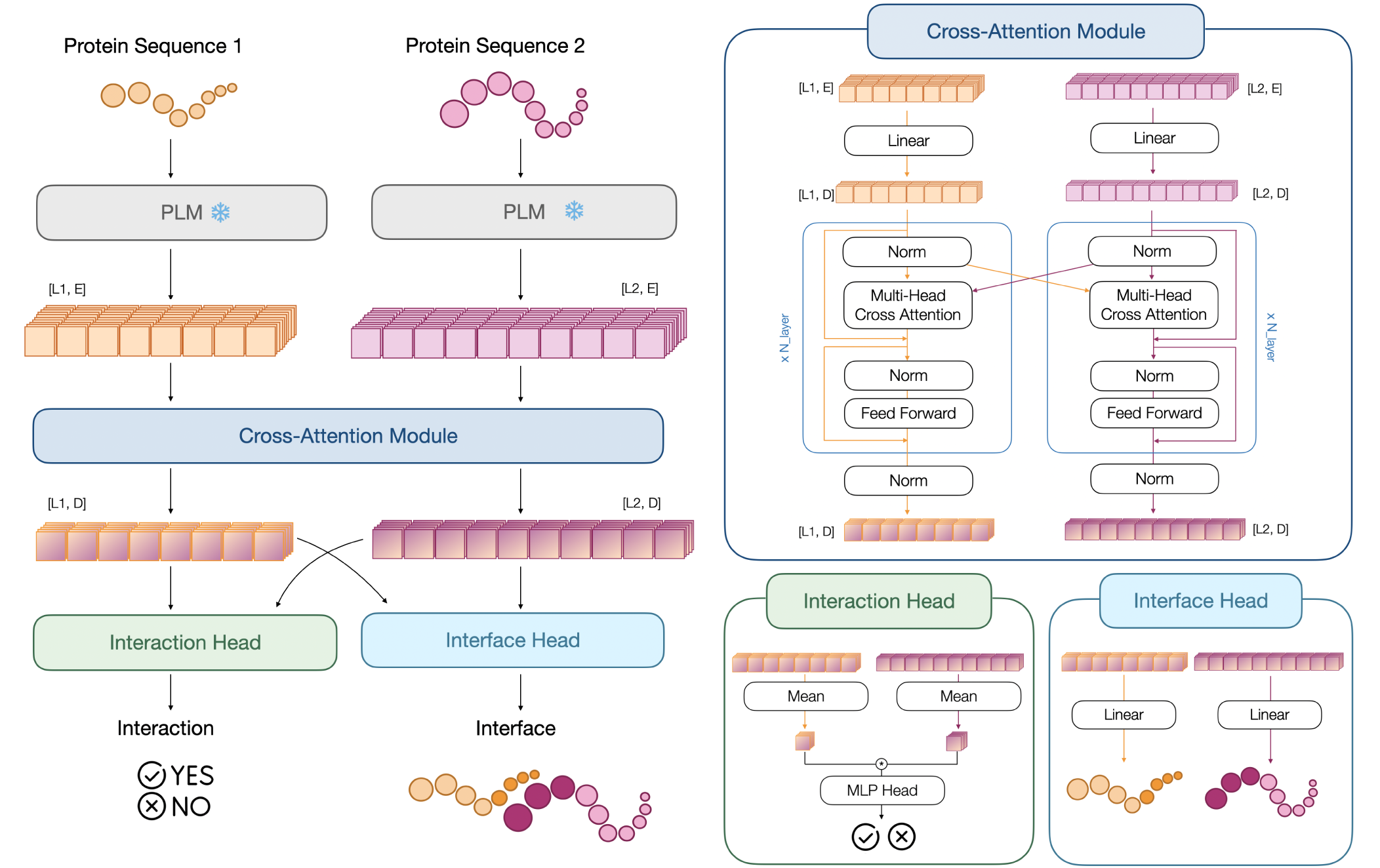
Overview of the X-PAIR multitask architecture. Two protein sequences are first encoded into embeddings using a pretrained PLM (Ankh). These representations are then processed through a cross-attention module that enables information exchange between the two sequences, capturing inter-protein dependencies. The resulting features are fed into two task-specific prediction heads: one for interaction prediction and one for residue-level interface prediction. The model can be trained either in a multitask setting, jointly optimizing both objectives, or in a single-task configuration focusing on just one task.

In the mammalian experiment, consistent with all other lineages, AUROC performance appears largely independent of evolutionary divergence time, remaining relatively stable even for species that are evolutionarily distant from the present. This indicates that the model does not exhibit a systematic degradation in predictive quality as evolutionary divergence time increases, suggesting that the underlying sequence embeddings retain informative features even in more remote regions of sequence space. Moreover, although performance is often expected to strongly depend on sequence similarity to the training set, our analysis reveals only a consistent but limited association (Fig. S7). Across most species, mean sequence identity shows a statistically significant monotonic relationship with AUROC, as measured by Spearman’s rank correlation coefficient *ρ*. However, this association remains modest: even in plants, *Archaea*, and *Fungi*, where *ρ >* 0.3, linear regression models yield *R*^2^ values below 18%, indicating that only a limited fraction of the variability in AUROC is captured by a linear relationship with sequence identity. In insects and mammals, the relationship is even more marginal (*ρ <* 0.3), while in viruses it is largely absent (*ρ ≈* 0.05). Overall, these results suggest that, although higher identity to the training set is associated with better performance, this relationship is limited, with most of the variability in AUROC likely reflecting additional factors, including functional conservation, local conformational properties, and the possibility that interaction interfaces retain features already represented in the training data even when sequence identity is low.

We then conducted a more fine-grained analysis focusing specifically on viral proteins, motivated by their markedly different evolutionary timescales: viral lineages evolve over much shorter time horizons, which could in principle lead to distinct generalisation behaviours compared to cellular organisms. Therefore, we analysed how different viral species vary with similarity to the training set, while also disentangling potential differences between intra-species and inter-species interaction settings. In particular, we first evaluated viral chains involved in within-species virus–virus interactions, observing consistent performance across viral groups (Fig. 5C). We then examined viral chains interacting with proteins from different mammalian hosts and *in vitro* systems, again finding broadly stable performance with no clear dependence on the interaction partner species (Fig. 5D). Across the 210 viral species represented in both analyses, intra-species interactions show a modest but statistically significant AUROC advantage over inter-species interactions (median paired difference in mean AUROC = +0.035; paired Wilcoxon signed-rank test, *p* = 2.3 *×* 10*^−^*^3^). Overall, this small but consistent effect suggests that, for viral proteins, within-species interactions are slightly easier to predict than cross-species ones.

## DISCUSSION

X-PAIR unifies interaction prediction and interface localisation within a single framework. Although biologically coupled, these tasks are not interchangeable: complex-structure predictors may generate plausible interfaces without reliably distinguishing partners from non-partners^17,18^, whereas binary PPI classifiers can identify interaction signals without resolving the residues that mediate binding^21–24^. By jointly predicting partner compatibility and partner-specific interfaces, X-PAIR connects network-level reconstruction with residue-level mechanistic information, while allowing both tasks to benefit from shared representations.

Individual examples in Fig. 3 illustrate that a high interaction score does not necessarily coincide with a clear interface prediction, and vice versa. Although such discrepancies may reflect model limitations, incomplete annotations or particularly difficult protein pairs, the large-scale evolutionary analyses provide stronger evidence that the two tasks rely, at least partly, on distinct biological signals. Interaction prediction declines gradually as the evolutionary distance between test sequences and the training distribution increases (Fig. 4), suggesting sensitivity to proteome composition, evolutionary history and similarity to the training distribution. Multispecies training further improves PPI prediction (Fig. 4), indicating that exposure to cross-species diversity helps the model learn broader interaction rules and reduces its dependence on species-specific biases. The magnitude of this improvement, relative to the limited increase in sequence similarity to the training set, suggests that the benefit arises from exposure to diverse proteomes rather than simply from the inclusion of closer homologues.

Interface localisation follows a different generalisation regime. Its performance remains comparatively stable across evolutionary distances and taxonomic groups (Fig. 5) and improves less markedly when additional species are included during training (Fig. 4). This behaviour is consistent with a stronger dependence on local physicochemical and evolutionary determinants of molecular recognition that are broadly conserved and encoded within protein sequences^40^. The contrasting evolutionary behaviours of the two tasks indicate that partner discrimination and interface localisation are complementary rather than simply different resolutions of the same prediction problem. These differences have practical implications for dataset design: multispecies training may be particularly important for partner discrimination, whereas interface localisation may rely more strongly on conserved local signals. An important practical question is therefore which combination of training species best balances evolutionary diversity, relevance to the target proteome and predictive accuracy.

The multitask formulation also introduces little additional computational cost. For 1,000*×*1,000 protein pairs, corresponding to one million candidate interactions, X-PAIR requires approximately 95 minutes for interaction prediction, 100 minutes for interface prediction and 105 minutes for both. Residue-level mechanistic information can therefore be obtained alongside interaction predictions with only a marginal increase in inference time, at an exceptionally high throughput of more than 150 protein pairs per second. Among the methods evaluated, PLM-interact and PIONEER were the best-performing task-specific baselines for PPI prediction and interface localisation, respectively. For the same number of protein pairs, PLM-interact would require more than 2 days for PPI prediction alone, whereas PIONEER would require more than 30 days for interface prediction alone. By contrast, X-PAIR completes both tasks jointly in under 2 hours. Together with its sequence-only design, this efficiency enables proteome-scale reconstruction of PPI networks in which predicted edges are annotated with the residues most likely to mediate each interaction.

Beyond its computational efficiency and multitask formulation, X-PAIR also achieves the strongest performance in single-task evaluations, with interaction prediction and partner-specific interface localisation assessed separately on two distinct benchmarks. For interaction prediction, X-PAIR slightly outperforms competing methods on the Gold Standard benchmark^29^, which was specifically designed to minimise sequence leakage. On the separate Xiong et al. benchmark for partner-specific interface localisation^4^, X-PAIR achieves the best overall performance among existing methods, with a particularly marked advantage in AUPRC (0.353) over PIONEER, the second-best-performing method (AUPRC = 0.256). Together, these results represent a substantial advance toward disentangling binding signals in proteins with multiple interaction partners^41^. The *ab initio* reconstruction of PPI networks together with partner-specific interfaces could reveal how protein surfaces are reused across different molecular contexts and help quantify their roles in molecular recognition, network organisation and protein dysfunction.

The value of partner-specific interface prediction is particularly apparent for proteins with multiple interaction partners, whose surfaces may be reused across different molecular contexts. Classical interface predictors identify regions with evolutionary, physicochemical or geometrical properties compatible with molecular recognition, but do not necessarily determine which partner binds a given surface^42–45^. Docking studies similarly show that the same surface region may accommodate both cognate partners and non-partners, such that low-energy docking solutions can reflect generic physical compatibility rather than a unique biological interaction^14,16,46,47^. Partner-specific interface prediction therefore provides information not only about whether a surface is interaction-compatible, but also about how that surface may be differentially used across partners.

This reuse of interaction-compatible surfaces is also related to protein-level “sociability” or “stickiness”^47,48^. Some proteins are broadly prone to forming molecular contacts and may interact with many candidate partners, whereas others display more selective interaction behaviour. These protein-specific tendencies can provide additional information for network reconstruction. Interaction indices that account for the behaviour of each protein across a set of potential partners have been shown to improve reconstruction confidence^16,47^. Future deep learning architectures could exploit this principle by reasoning not only over isolated protein pairs, but also over the interaction profile of each protein across a broader distribution of candidate partners within a proteome.

Proteome-scale inference also creates opportunities to model how mutations and natural sequence variation affect both interaction probabilities and partner-specific interfaces. Even subtle substitutions, particularly at interface residues, may alter binding affinity or disrupt an interaction. Predicting these effects across a proteome could reveal mutation-induced network rewiring and support the reconstruction of condition- or individual-specific interactomes. A key future direction is therefore to make proteome-scale PPI predictions quantitatively sensitive to sequence perturbations, enabling the reconstruction of mutation-specific interaction networks that reveal the mechanistic basis and functional consequences of network rewiring.

In summary, X-PAIR establishes sequence-based multitask learning as a scalable framework for PPI reconstruction and partner-specific interface prediction. By combining proteome-wide efficiency, cross-species generalisation and residue-level mechanistic information, it provides a foundation for models that capture the dynamic, evolutionary and context-dependent organisation of PPI networks.

## METHODS

### Data processing

A central contribution of this work is the construction of curated datasets for the prediction of PPIs and residue-level interfaces. These datasets were designed not only to train and evaluate X-PAIR, particularly in its multitask setting, but also to provide reusable benchmarks for future models addressing the coupled problems of interaction prediction and interface localisation.

To this end, interface data were collected from PPI3D^49^, which provides protein pairs derived from PDB structures together with residue-level annotations of the corresponding binding interfaces. Interaction data were obtained from STRING v12.0^50^, a large-scale database containing physical PPIs across multiple species. Importantly, interface data from PPI3D may include cross-species interactions, whereas STRING-derived interaction data are restricted to PPIs between proteins from the same species. The two data sources were processed through distinct pipelines, both aimed at retaining biologically meaningful, high-quality, and non-redundant protein pairs. Moreover, interaction data were further processed to generate high-quality negative pairs following the procedure introduced in SENSE-PPI^3^. In this section, we describe in detail the two different pipelines used to process and generate the interface data derived from PPI3D and the interaction data derived from STRING. These pipelines were then applied to build several training and test dataset collections, as described in the Datasets section of the Methods.

### Pipeline to process interface data

The interface data were derived from PPI3D^49^, a curated resource of interacting protein pairs with experimentally resolved three-dimensional structures. PPI3D aggregates biological assemblies of all non-NMR structures in PDB and provides residue-level interface annotations together with structural and sequence metadata. Residue interfaces in PPI3D are defined using Voronoi tessellation, a geometric partitioning of three-dimensional space in which each atom of a protein is assigned a region containing all points closer to it than to any other atom. This defines geometric contacts between atoms that can then be aggregated at the residue level. In this way, the PPI interface is defined as the set of residue–residue contacts occurring between different chains. All PPI pairs with PDB resolution *<* 3.5 Å and release dates spanning from 1973-01-01 to 2025-10-08 were retrieved from PPI3D, together with the structural annotations of the associated interfaces. These structural interface annotations were then, whenever possible, remapped onto the corresponding protein sequences. In 25 cases this remapping was not possible, and the corresponding interaction pairs were therefore discarded. The resulting data comprised 786,533 interaction pairs. Several quality-control filters were then applied to ensure data reliability and to minimise biases that could adversely affect model training. In particular, for each PDB only the first biological assembly was retained to avoid inconsistencies arising from alternative quaternary-structure reconstructions. Proteins with more than 30% residues lacking atomic coordinates or more than 30% ambiguous residues (“X” residues) were excluded, as such entries were considered not to provide sufficiently reliable structural or sequence information to accurately describe interaction interfaces. Additional filters removed sequences shorter than 50 residues or longer than 2,000 residues, both to optimise computational efficiency and to avoid biases associated with unstable fragments or extremely large proteins. Interaction pairs in which one partner exhibited more than 70% of residues labeled as interacting were also discarded, since these cases typically correspond to crystallographic contacts or artefactual interfaces rather than biologically meaningful interactions. Furthermore, pairs with buried surface area *<* 300 Å ^2^ were removed to retain only sufficiently large and biologically relevant interfaces. For homodimeric complexes, residue-level interfaces of the two chains were unified so that both shared identical interface definitions, ensuring annotation consistency. After these filtering steps, the final curated database contained 466,861 interaction pairs composed of 83,367 unique sequences from 105,252 PDB structural complexes. From these curated interface data, we generated multiple downstream datasets tailored to the specific requirements of each experiment and model-training procedure, as described in the Datasets section of the Methods. In particular, each dataset was constructed by retaining only protein pairs belonging to the species or taxonomic group relevant to the corresponding experiment. Duplicate sequence pairs were then consolidated by merging their residue-level interfaces through set union, enabling the model to learn the full set of experimentally observed interaction residues across alternative conformations of the same protein pair. Finally, redundancy reduction was performed using MMseqs2^38^ clustering at 90% sequence identity, retaining within each cluster pair only the representative interaction pair with the maximum combined sequence length. A relatively high identity threshold was intentionally maintained to preserve subtle sequence variations that may influence residue-level interface formation while avoiding excessive loss of already limited structural interface data. This choice allows the model to learn to discriminate interaction interfaces even among highly similar sequences without substantially reducing dataset size.

### Pipeline to process interaction data

PPI data were generated using the create_dataset function from the SENSE-PPI package^3^, which implements a standardized pipeline for constructing positive and negative interaction pairs while limiting the inclusion of false negatives through a “neighbouring-exclusion” strategy. In particular, interaction data are extracted from physical PPIs reported in STRING v12.0^50^. Given a species of interest, the function retains only interactions between proteins belonging to that species, with a combined confidence score above a user-defined threshold, and filters proteins according to a specified sequence-length range. To reduce redundancy and mitigate biases due to highly similar sequences, the function clusters proteins using MMseqs2 at 40% sequence identity and selects a single representative interaction for each pair of interacting clusters. Negative pairs are also generated within create_dataset, using the “neighbouring-exclusion” procedure introduced with SENSE-PPI^3^. Specifically, a protein pair A–B is considered as a valid negative example only if (i) no interaction between A and B is reported in STRING, and (ii) protein B does not belong to the same sequence cluster as any known interactor of A. This strategy ensures that negative examples are defined not only by the absence of annotated interactions but also by minimising homology-driven biases within shared sequence clusters. Finally, the procedure fixes the ratio of positive to negative protein pairs at 1:10, in order to approximate the natural imbalance of interaction frequencies observed in biological systems. Data for multiple species were generated independently using the create_dataset function of SENSE-PPI, by applying the following criteria: (i) a confidence score threshold of 500, and (ii) protein sequence lengths restricted to the range of 50 to 2,000 amino acids, in order to optimise computational efficiency and avoid biases associated with extremely short or long sequences. As described in the Datasets section, different subsets of species were used for model training, while others were reserved for evaluation. For training species, all positive interaction pairs satisfying criteria (i) and (ii) were included to maximise interaction diversity. In contrast, for evaluation species, the same criteria (i) and (ii) were applied, together with an additional constraint limiting the data to at most 5,000 positive interaction pairs (--max_positive_pairs), yielding balanced and comparable test sets across organisms.

### Datasets

The pipelines described in the previous paragraph for processing the interface data derived from PPI3D and the interaction data derived from STRING were used to generate several training and test dataset collections. These datasets were designed to support different experimental settings and to evaluate distinct aspects of model performance. In particular, four dataset collections were generated: X-fair, X-human, X-multispecies, and X-taxonomic-interface.

- **X-fair**: this dataset collection was designed to reduce information leakage between training, validation, and test sets. Given the limited availability of residue-level interface annotations, all biologically meaningful interface samples obtained from the procedure described above were retained. The STRING-derived interaction data were instead restricted to four representative species: *Homo sapiens*, *Gallus gallus*, *Saccharomyces cerevisiae*, and *Drosophila melanogaster*. To define homology-aware training, validation, and test splits, proteins from both the interface and interaction datasets were first merged and then clustered using MMseqs2 at 30% sequence identity and 80% sequence coverage. Splits were then assigned at the cluster-pair level, ensuring that each pair of clusters appeared exclusively in a single split. This strategy prevents homologous protein pairs from being distributed across different splits, thereby reducing homology-based leakage and enabling a stringent evaluation of generalisation performance across both tasks. The resulting split comprised 1,724,263/215,533/215,533 interaction pairs and 41,938/5,242/5,242 interface samples for the training, validation, and test sets, respectively.
- **X-human**: this dataset collection was designed to evaluate model generalisation from an evolutionary perspective, by training and validating on human proteins and testing on proteins from other species. For both interaction and interface prediction, the training and validation sets were restricted to protein pairs in which both partners belong to *Homo sapiens*. The training set comprised 879,318 human interaction pairs and 7,997 human interface samples, while the validation set included 97,702 human interaction pairs and 889 human interface samples. Generalisation was then evaluated on species-specific test sets from *Escherichia coli*, *Arabidopsis thaliana*, *Caenorhabditis elegans*, *Bombyx mori*, *Danio rerio*, *Oryctolagus cuniculus*, *Sus scrofa*, and *Bos taurus*, with both proteins in each test pair belonging to the same species. The sizes of the corresponding test datasets are reported in Table S2.
- **X-multispecies**: this dataset collection was also designed to assess evolutionary generalisation, but using a broader multispecies training set. In this case, both interaction and interface training and validation data were restricted to intra-species protein pairs from four species: *Homo sapiens*, *Gallus gallus*, *Saccharomyces cerevisiae*, and *Drosophila melanogaster*. The combined training set comprised 1,939,797 interaction pairs and 9,862 interface samples, whereas the validation set included 215,533 interaction pairs and 1,096 interface samples. The test species were the same as those used for X-human, allowing a direct comparison between human-only and multispecies training strategies.
- **X-taxonomic-interface**: this dataset collection comprised exclusively interface data and was used to evaluate the single-task version of X-PAIR across different evolutionary scales. For each taxonomic group of interest (e.g. mammals), the training set was restricted to pairs in which neither partner belonged to the target group, while the test set included all pairs with at least one protein from that group. Interface prediction performance was then evaluated on all proteins belonging to the target group, regardless of the species of their interaction partners. This procedure was applied to the following taxonomic groups: *Mammalia*, *Viridiplantae*, *Viruses*, *Archaea*, *Fungi*, and *Insecta*. The sizes of the corresponding datasets are reported in Supplementary Table S3.

Furthermore, to evaluate the performance of the single-task version of X-PAIR, the model was trained and tested on three well-established benchmark datasets. This allows the model to be assessed in a fair and standardised setting and facilitates comparison with existing methods. For this evaluation, we used three available benchmarks: the PIONEER dataset for interface benchmarking^4^, the cross-species dataset for interaction benchmarking^21^, and the Gold Standard dataset for interaction benchmarking^29^. A detailed description of these datasets is provided in the Supplementary Material, as well as in their original publications. All training, validation, and test splits used in this work are made available online to ensure reproducibility and to facilitate reuse of the datasets for model training and benchmarking by other researchers.

### Model architecture

X-PAIR is a multitask neural model designed to jointly predict PPIs and residue-level interaction interfaces from sequence representations. The model follows a Siamese architecture, ensuring commutativity with respect to the order of the input proteins.

The model takes as input two protein sequences of variable lengths *L*_1_ and *L*_2_, which are padded within each mini-batch to a common length. Each input sequence is first encoded into residue-level representations using a pretrained protein language model (PLM). Although X-PAIR can use different PLM encoders, including ESM-2^51^ and ESM C^52^, all experiments in this study use Ankh-large (1.15B parameters)^53^, which provided the best trade-off between predictive performance and model size (see the Protein language model selection section). The protein embeddings are then projected into a shared latent space of dimension *D* = 128 through a linear layer, reducing computational cost while preserving expressive power. The projected representations are processed by a stack of two bidirectional cross-transformer layers^54^. Each layer follows a pre-normalization scheme^55^ and is built around a 2-head rotary position-encoded (RoPE)^56^ cross-attention mechanism, complemented by residual connections and a position-wise feed-forward network. RoPE injects relative positional information directly into the query–key interactions through position-dependent rotations, enabling the model to capture distance-aware relationships between residues and to generalise across variable sequence lengths. Within each layer, cross-attention is applied symmetrically in both directions, allowing explicit exchange of inter-chain information while preserving sequence-specific representations. A final layer normalization is applied before the prediction heads to improve numerical stability.

On top of this shared backbone, X-PAIR includes two task-specific prediction heads. The interaction head aggregates residue-level representations via masked mean pooling for each protein, combines the resulting protein-level embeddings through element-wise multiplication, and predicts a pairwise interaction logit using a multilayer perceptron composed of Linear(128 *→* 64), ReLU activation, dropout, and Linear(64 *→* 1). In parallel, the interface head performs residue-level classification by applying a shared linear layer to each token representation, consisting of dropout followed by Linear(128 *→* 1), producing per-residue interface logits for both proteins. In total, the model comprises only 469K trainable parameters (with 8.3K parameters in the interaction head and 129 parameters in the interface head), in addition to 1.15B pretrained parameters inherited from the Ankh-large backbone.

### Implementation details

Training supports interaction-only, interface-only, and joint multitask optimization. In the multitask setting, at each optimization step, one task-specific dataloader is selected with probability proportional to its number of batches, and a mini-batch is sampled from the selected dataloader. The corresponding task-specific loss is then computed and backpropagated, and model parameters are updated exclusively based on the selected task loss. For interaction batches, the interaction loss is weighted by a factor of 2.0. This value was selected on the validation set as it provided the best overall trade-off between interaction and interface prediction performance. Across all training configurations, the interaction loss is defined as a protein-pair-level binary cross-entropy (BCE), whereas the interface loss is a mask-aware token-level BCE. To address class imbalance in interface annotations, a positive-class weight of 5 is applied, reflecting the approximately 17% prevalence of interface residues. Furthermore, any residues labeled X, indicating missing interface annotations, are explicitly masked during training and evaluation. These residues do not contribute to the interface loss computation and are excluded from all interface-related evaluation metrics.

X-PAIR is implemented in PyTorch using PyTorch Lightning. A dropout rate of 0.3 was applied across all transformer layers and prediction heads. Models were trained using AdamW (*lr* = 1 *×* 10*^−^*^4^) with a cosine learning-rate schedule and linear warmup. Training was performed for a maximum number of epochs depending on the size of the training dataset, with early stopping (patience = 3) based on validation loss. In the multitask setting, we used a batch size of 32 for interaction prediction and 1 for interface prediction. A complete summary of the hyperparameters used in each experiment is reported in Table S4. For all experiments, the random seed was fixed to 123, and the best checkpoint was selected based on the minimum global validation loss. All experiments were conducted on a single NVIDIA A100 (80 GB) GPU, except for the single-task interface prediction experiments across different taxonomic groups, which were performed on a single NVIDIA V100 GPU (32 GB).

### Protein language model selection

To identify the most suitable sequence encoder, we evaluated X-PAIR multitask models on the X-fair dataset using different pretrained PLMs. We considered models of different sizes, including ESM-2 650M and 3B^51^, ESM C 300M and 600M^52^, and Ankh 450M and 1.15B^53^. As shown in Fig. S1, Ankh 1.15B achieved the best performance for interface residue prediction, with the highest AUROC and AUPRC among the tested models. For PPI prediction, ESM-2 3B obtained the highest AUROC and AUPRC, although Ankh 1.15B performed very closely while using fewer parameters. Based on this trade-off between predictive performance and model size, we selected Ankh 1.15B as the backbone for the final X-PAIR model. Ankh is a general-purpose PLM based on a T5-like^57^ encoder–decoder Transformer architecture and optimised specifically for protein sequences. It was pretrained on UniRef50^58^ using a masking objective and achieved competitive performance across a broad range of protein structure and function prediction tasks, while requiring substantially lower computational resources than larger PLMs.

### MMseqs2

MMseqs2^38^ is a fast and sensitive tool for large-scale sequence similarity search and clustering of biological sequences. During dataset construction, it was used to cluster sequences with the *cluster* module using --alignment-mode 3, --cov-mode 1, and a coverage threshold of 0.8. Each protein pair was assigned to a cluster pair, and redundancy was reduced using minimum sequence identity thresholds of 0.9 for interface prediction and 0.4 for interaction prediction. The same framework was used to define homology-aware splits with a sequence identity threshold of 0.3 for both tasks. MMseqs2 was also used in the analysis phase to compute sequence identity between training and test proteins via the *search* module. Sequence identity was defined as the fraction of identical residues over all aligned positions, including gap-containing columns (--alignment-mode 3). A high-sensitivity configuration (-s 7.5) was used to ensure robust detection of homologous sequences. For each test protein, the maximum identity to any training protein was retained, with unmatched proteins assigned a value of 0. Species-level estimates were obtained by averaging these values across all proteins from each species.

### TimeTree

TimeTree^39^ is a database of species divergence times. In this study, it was used to reconstruct the phylogenetic tree and obtain the evolutionary distances shown in Fig. 4, as well as the divergence times reported in Fig. 5. For the latter analysis, each species was assigned the divergence time from its closest sister lineage available in TimeTree, used as a proxy for the time elapsed since it became a distinct evolutionary lineage.

### Decision threshold

Unless otherwise stated, all performance metrics were computed using a fixed decision threshold of 0.5 for both interaction and interface prediction, ensuring consistency across experimental settings. Nevertheless, model predictions can be further optimised by calibrating task-specific decision thresholds on validation data. In the X-fair training setting, the thresholds that maximised the validation F1 score were 0.40 for interaction prediction, yielding an F1 score of 0.82, and 0.66 for interface prediction, yielding an F1 score of 0.54. Thus, while a threshold of 0.5 was retained for standardised reporting, validation-calibrated thresholds may be preferable for downstream applications requiring optimal F1 performance.

## RESOURCE AVAILABILITY

### Code availability

X-PAIR is available at http://gitlab.lcqb.upmc.fr/srescalli/X-PAIR.git under the CC-BY-NC-SA 4.0 License. The repository contains the complete source code required to reproduce the analyses presented in this study, including the trained model checkpoints. The datasets generated for training and evaluating the models are publicly available on Zenodo at https://doi.org/10.5281/zenodo.21457017. Together, these resources enable full replication of the reported experiments and facilitate further benchmarking and evaluation by the community.

Links to the online tools used for structural visualization and phylogenetic analyses are provided below:

- PyMOL: https://pymol.org
- TimeTree: https://timetree.org

## Data availability

The benchmark datasets used for evaluation are derived from publicly available resources. The cross-species interaction dataset^21^ is available at https://zenodo.org/records/5140612, the Gold Standard interaction dataset^29^ at https://doi.org/10.6084/m9.figshare.21591618. v3, and the PIONEER interface dataset^4^ at https://static-content.springer.com/esm/art%3A10.1038%2Fs41587-024-02428-4/MediaObjects/41587_2024_2428_MOESM4_ESM.xlsx.

The datasets generated in this study were constructed from publicly available resources. Specifically, protein interaction information from the STRING database (https://string-db.org/) and structural interface annotations from PPI3D (https://bioinformatics.lt/ppi3d/ clusters) were used to generate the training and evaluation datasets described in this study. These datasets are publicly available on Zenodo at https://doi.org/10.5281/zenodo.21457017.

## ACKNOWLEDGMENTS

This work was performed using HPC resources from GENCI–IDRIS and Sorbonne University GPU clusters from SCAI and LIP6 laboratory (UMR 7606, Sorbonne University-CNRS). Financial support from PostGenAI@Paris within the framework of France 2030 ANR-23-IACL-0007 (AC), ANR ALLEGRO ANR-23-CE11-0006 (AC), and Institut Universitaire de France (AC).

## AUTHOR CONTRIBUTIONS

Conceptualization, S.R. and A.C.; methodology, S.R. and A.C.; software implementation, S.R.; data analysis, S.R. and A.C.; writing & editing, S.R and A.C.; funding acquisition, A.C.; supervision, A.C.

## Supplementary Information

### Supplementary Text

#### Benchmark datasets

##### Cross-species dataset for interaction benchmarking

The cross-species PPI benchmark introduced by Sledzieski et al. ^21^ consists of human interaction data for training and validation, together with independent test sets from five additional organisms: *M. musculus*, *D. melanogaster*, *C. elegans*, *S. cerevisiae*, and *E. coli*, all derived from STRING v11^59^. Positive samples correspond to experimentally supported physical interactions, whereas negative examples are generated by randomly pairing proteins without reported interactions. The resulting datasets exhibit a positive-to-negative ratio of approximately 1:10, reflecting the inherent sparsity of true PPIs in biological networks. Protein lengths range from 50 to 800 amino acids, and interaction pairs are clustered at 40% sequence identity using CD-HIT to reduce redundancy. The human training split includes 421,792 protein pairs, and the validation set comprises 52,725 pairs. Each non-human species test set contains 55,000 interactions, except for *E. coli*, which includes 22,000 due to the lower number of experimentally supported PPIs available in STRING.

##### Gold Standard dataset for interaction benchmarking

The leakage-free interaction benchmarking dataset constructed by Bernett et al. ^29^ is based on experimentally supported human PPIs curated from HIPPIE v2.3^60^. Negative interaction pairs were generated through random sampling while preserving node degree distributions in expectation, resulting in a balanced dataset with a 1:1 ratio of positive to negative interactions. The dataset was partitioned using the KaHIP graph partitioning strategy^61^ to ensure separation between splits, and minimisation of sequence similarity among training, validation and test datasets. Furthermore, redundancy reduction was applied using CD-HIT^62^ at a 40% sequence identity threshold, ensuring that proteins are pairwise dissimilar both across and within splits. This stringent design compels models to extract interaction-relevant features beyond simple sequence similarity. For computational reasons, in our experiments protein pairs containing sequences longer than 2,000 amino acids were excluded from the training and validation splits, while evaluation was conducted on the full test benchmark, resulting in 146,089 interactions for training, 56,051 for validation and 52,048 for testing.

##### PIONEER dataset for interface benchmarking

The interface benchmark introduced in PIONEER^4^ comprises partner-specific residue-level interface annotations derived from experimentally determined PPIs with available co-crystal structures across eight species (*H. sapiens*, *A. thaliana*, *S. cerevisiae*, *D. melanogaster*, *C. elegans*, *M. musculus*, *S. pombe* and *E. coli*). Interface residues were identified using NACCESS^63^ by measuring changes in solvent-accessible surface area (SASA) upon complex formation. A residue was labeled as interfacial if it was surface-exposed (*≥* 15% relative SASA) and exhibited a decrease of at least 1Å ^2^ in the complex. All available co-crystal structures for a given interaction were considered, and a residue was marked as interface if it satisfied the criterion in at least one structure. Only interactions with structural coverage of at least 30% of UniProt residues for both proteins were retained. To prevent information leakage, no homologous interactions or repeated proteins were shared across splits. Homology was assessed using three iterations of PSI-BLAST with an E-value cutoff of 0.001, resulting in 2,615 training interactions, 400 validation interactions, and 400 independent test interactions with sequence length between 20 and 4,128. Approximately 12% of residues were annotated as interface, while 22% lacked interface annotations. Residues with missing interface labels were excluded from both loss computation and performance evaluation. In our experiments, protein pairs containing sequences longer than 2,000 amino acids were removed from the training and validation splits to maintain computational tractability, resulting in 2,599 pairs for training, 400 for validation and 400 for testing.

## Supplementary Tables

**Table S1:**
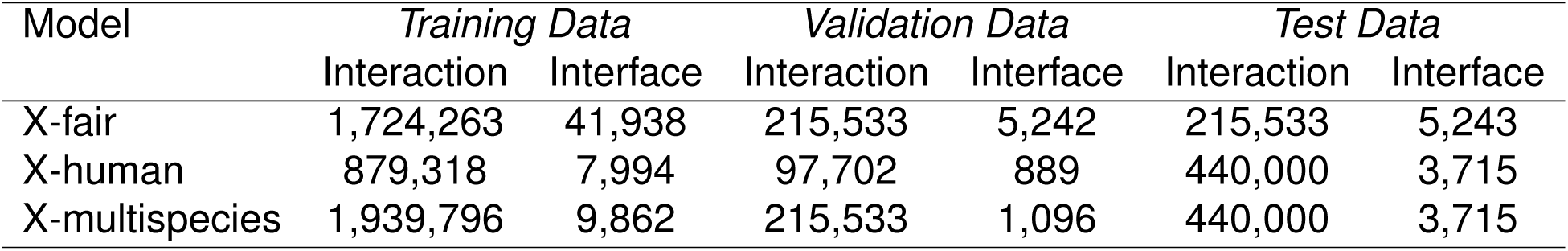
Sizes of the datasets generated in this study and used for training X-PAIR under different experimental settings, reported separately for interaction prediction and interface prediction tasks.

**Table S2:** Number of protein pairs for each species in the test sets of the X-human and X-multispecies datasets.

| Test species | Scientific Name | Taxon ID |  | Test Data |  |
| --- | --- | --- | --- | --- | --- |
|  |  | Interaction | Interface | Interaction | Interface |
| E. coli | *Escherichia coli* | 511145 | 562 | 55,000 | 1,484 |
| Thale cress | *Arabidopsis thaliana* | 3702 | 3702 | 55,000 | 614 |
| Worm | *Caenorhabditis elegans* | 6239 | 6239 | 55,000 | 203 |
| Silkworm | *Bombyx mori* | 7091 | 7091 | 55,000 | 26 |
| Zebrafish | *Danio rerio* | 7955 | 7955 | 55,000 | 131 |
| Rabbit | *Oryctolagus cuniculus* | 9986 | 9986 | 55,000 | 210 |
| Pig | *Sus scrofa* | 9823 | 9823 | 55,000 | 513 |
| Cow | *Bos taurus* | 9913 | 9913 | 55,000 | 534 |
| Total |  |  |  | 440,000 | 3,715 |

**Table S3:** Sizes of the X-taxonomic-interface datasets for each held-out taxonomic group.

| Tested taxonomic group | Train | Validation | Test |
| --- | --- | --- | --- |
| *Archaea* | 45,459 | 5,052 | 1,944 |
| *Fungi* | 43,399 | 4,823 | 4,203 |
| *Insecta* | 46,610 | 5,179 | 659 |
| *Mammalia* | 32,477 | 3,609 | 16,576 |
| *Viridiplantae* | 44,525 | 4,948 | 2,957 |
| *Viruses* | 43,038 | 4,783 | 4,593 |

**Table S4:** Hyperparameters used for training X-PAIR across different benchmarks and datasets.

| Task | Training dataset | Max Epochs | Warmup steps | Interaction batch size | Interface batch size |
| --- | --- | --- | --- | --- | --- |
| Interaction | Gold Standard benchmark<sup>29</sup> | 6 | 1.000 | 32 | - |
| Interaction | Cross-species benchmark<sup>21</sup> | 8 | 3.000 | 32 | - |
| Interface | PIONEER benchmark<sup>4</sup> | 8 | 20 | - | 32 |
| Multi-task | X-fair | 10 | 20.000 | 32 | 1 |
| Multi-task | X-human | 10 | 20.000 | 32 | 1 |
| Multi-task | X-multispecies | 10 | 20.000 | 32 | 1 |

**Table S5:** Performance of X-PAIR on the X-fair dataset for both interaction and interface prediction tasks. Results are reported for both the multitask and single-task settings on the corresponding datasets.

| Task | Setting | AUROC | AUPRC | Recall | Precision | F1 | MCC |
| --- | --- | --- | --- | --- | --- | --- | --- |
| Interaction | multitask | 0.97 | 0.87 | 0.76 | 0.87 | 0.81 | 0.80 |
|  | Single-task | 0.97 | 0.89 | 0.77 | 0.89 | 0.83 | 0.82 |
| Interface | multitask | 0.83 | 0.56 | 0.74 | 0.37 | 0.49 | 0.39 |
|  | Single-task | 0.84 | 0.57 | 0.75 | 0.37 | 0.49 | 0.39 |

**Table S6:** Evaluation of residue-level interface prediction using top-*k* precision (P@k) and recall (R@k). These metrics are computed by ranking residues in each complex by predicted interface probability, measuring precision and recall within the top-k selected residues, and then averaging the resulting values across complexes as described by Morehead et all^64^. Metrics are reported for *k* = 20*, L/*10*, L/*5, where *L* denotes the sum of the lengths of the two proteins.

| Model | P@20 | P@L/10 | P@L/5 | R@20 | R@L/10 | R@L/5 |
| --- | --- | --- | --- | --- | --- | --- |
| multitask | 0.57 | 0.51 | 0.44 | 0.15 | 0.30 | 0.49 |
| Interface-only | 0.58 | 0.51 | 0.44 | 0.15 | 0.30 | 0.50 |

**Table S7:** Training and inference times of X-PAIR on the X-fair dataset in the multitask, interaction-only, and interface-only settings. Reported values include the total training time, the time spent generating training and validation embeddings, the total test time, and the time required to generate test embeddings. All embeddings were generated using the pretrained Ankh-large model.

| Model | *Time for Training (emb. generation)* | *Time for Testing (emb. generation)* |
| --- | --- | --- |
| multitask | 52h (1h) | 59m (33m) |
| Interaction-only | 43h (28m) | 54m (28m) |
| Interface-only | 4h (32m) | 6m (5m) |

## Supplementary Figures

**Figure S1:**
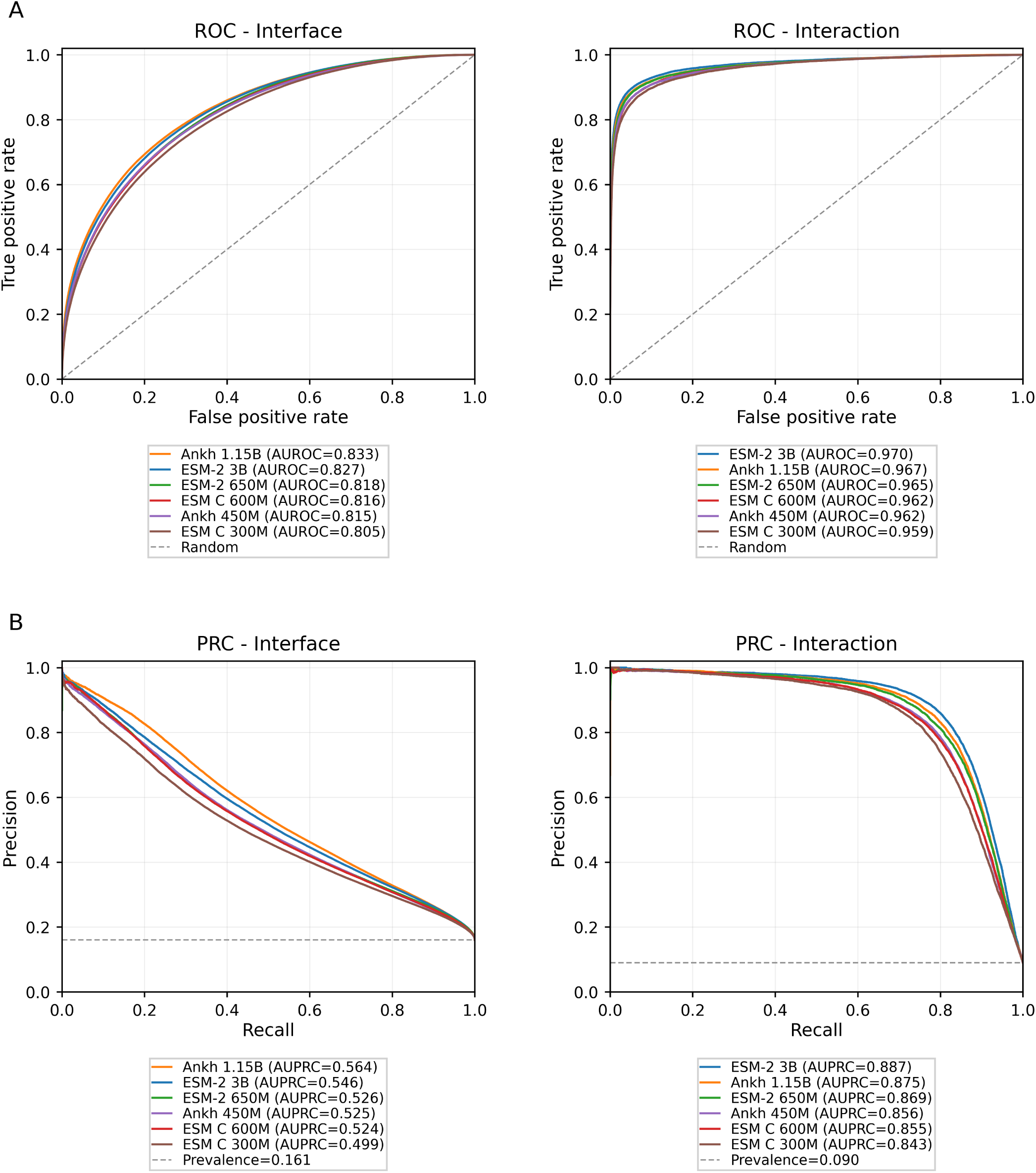
Performance of multitask X-PAIR models using different pretrained PLMs. (A) ROC curves for interface residue prediction and PPI prediction. (B) Precision–recall curves for the same two tasks. The curves compare multitask X-PAIR models trained on the X-fair dataset using representations from different pretrained PLMs. Legends report AUROC for ROC curves and AUPRC for precision–recall curves. AUPRC was computed as average precision. Random and prevalence baselines are shown as dashed grey lines in the ROC and precision–recall plots, respectively.

**Figure S2:**
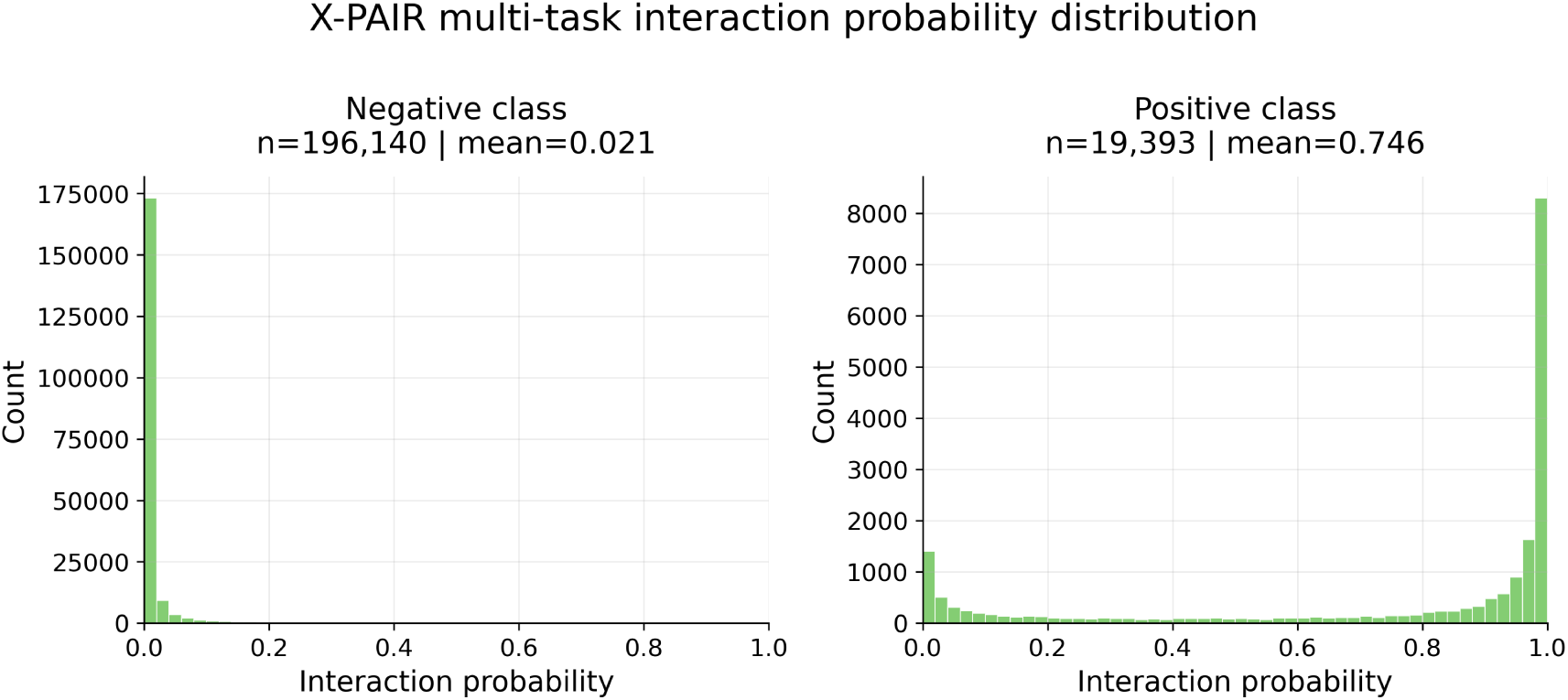
Distribution of interaction probabilities predicted by X-PAIR multitask on the X-fair interaction test set. Histograms show predicted probabilities for annotated positive and negative pairs.

**Figure S3:**
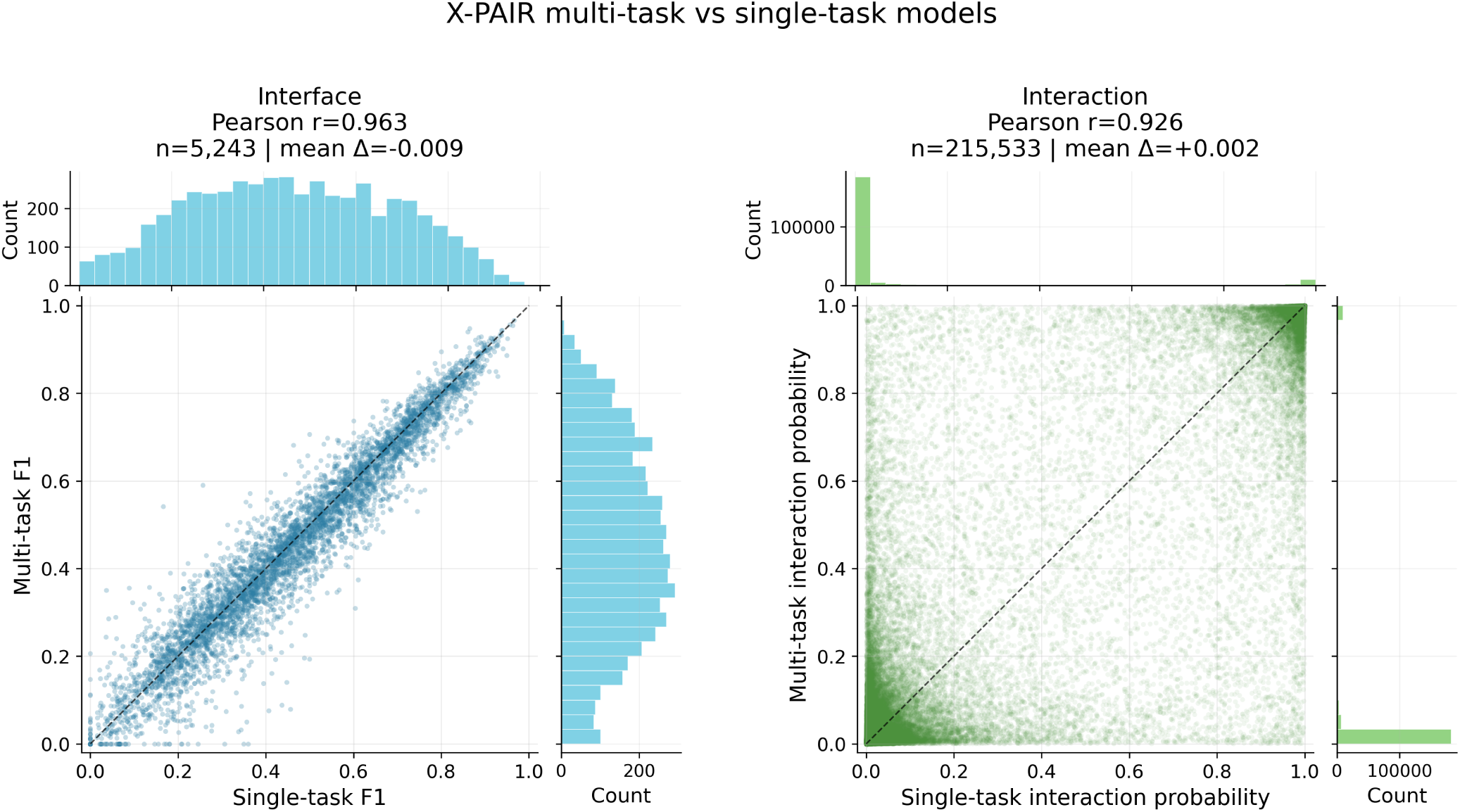
Comparison between single-task and multitask X-PAIR models on the X-fair dataset. Scatter plots compare the outputs of the single-task and multitask X-PAIR models on the same X-fair test examples for both the interaction and interface tasks. Interface predictions are summarised using the pair-level F1 score, whereas interaction predictions are compared using the predicted interaction probability. Marginal histograms show the corresponding output distributions. The Pearson correlation reports the linear agreement between single-task and multitask outputs. The number of test examples and the mean difference between the two models are also reported for both tasks.

**Figure S4:**
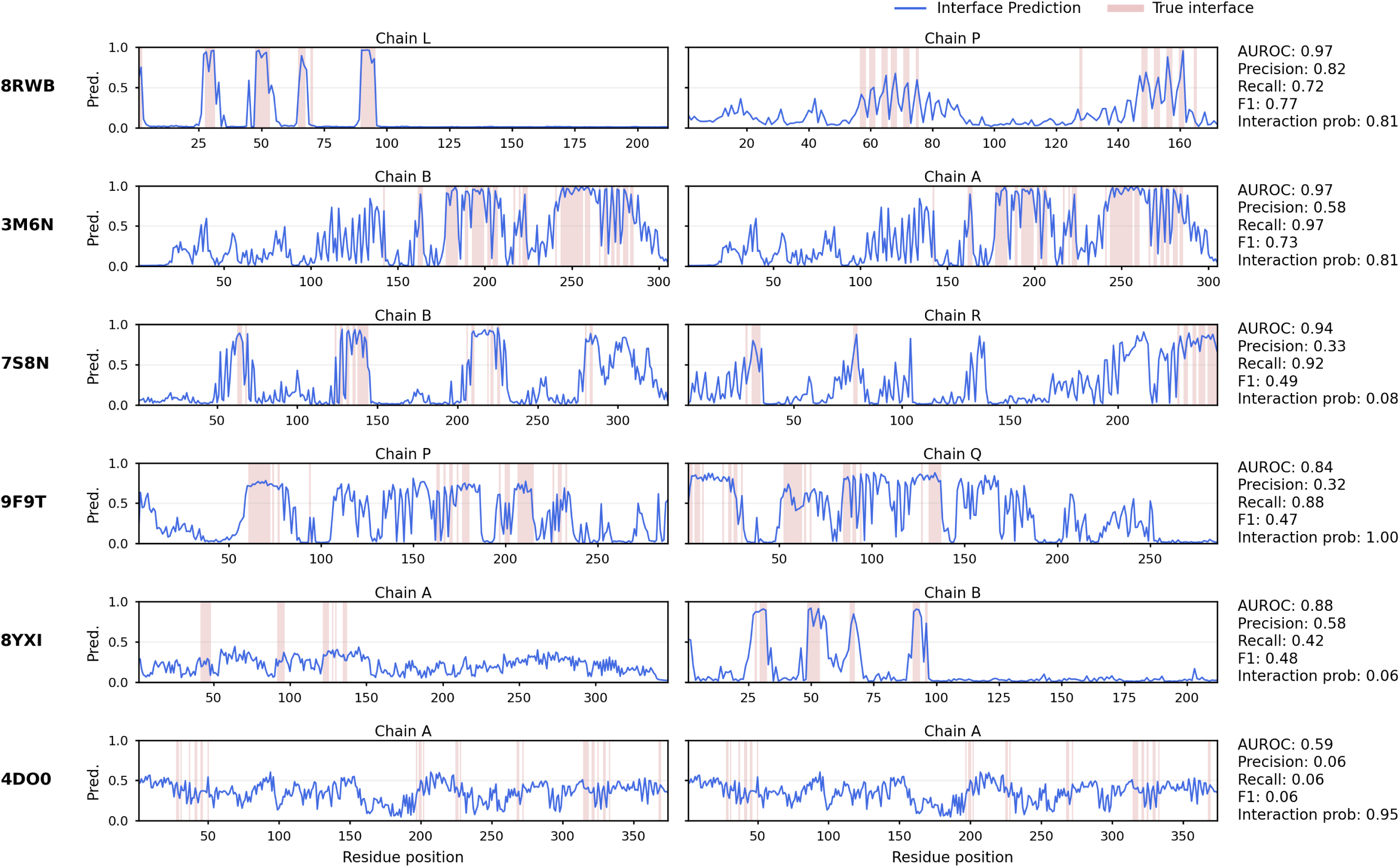
X-PAIR interface predictions for each chain pair of the selected PDB examples shown in Fig. 3. Blue profiles indicate the predicted probability that a residue in a chain belongs to the interface, while red bands mark the corresponding ground-truth interface residues. For each chain pair, the interface-level performance metrics for the pair are reported on the right, together with the interaction probability predicted by X-PAIR.

**Figure S5:**
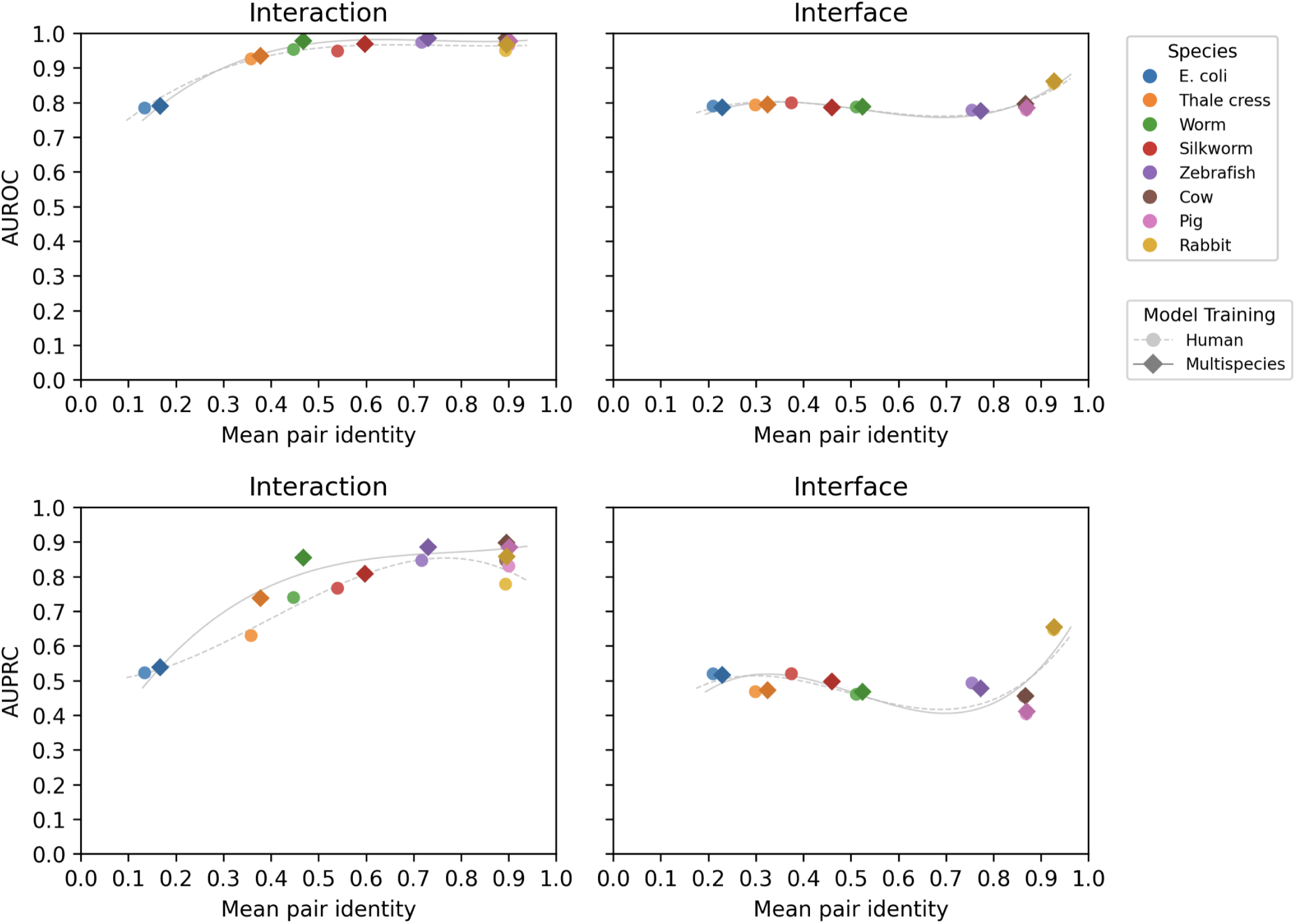
Cross-species generalisation of interaction and interface prediction. AUROC and AUPRC shown versus mean pairwise sequence identity between test proteins and the training data, across multiple species. Results are reported for interaction prediction at the pair level and interface prediction at the residue level, comparing X-PAIR trained exclusively on human proteins (X-human dataset) with X-PAIR trained on human, chicken, fly, and yeast proteins (X-multispecies dataset).

**Figure S6:**
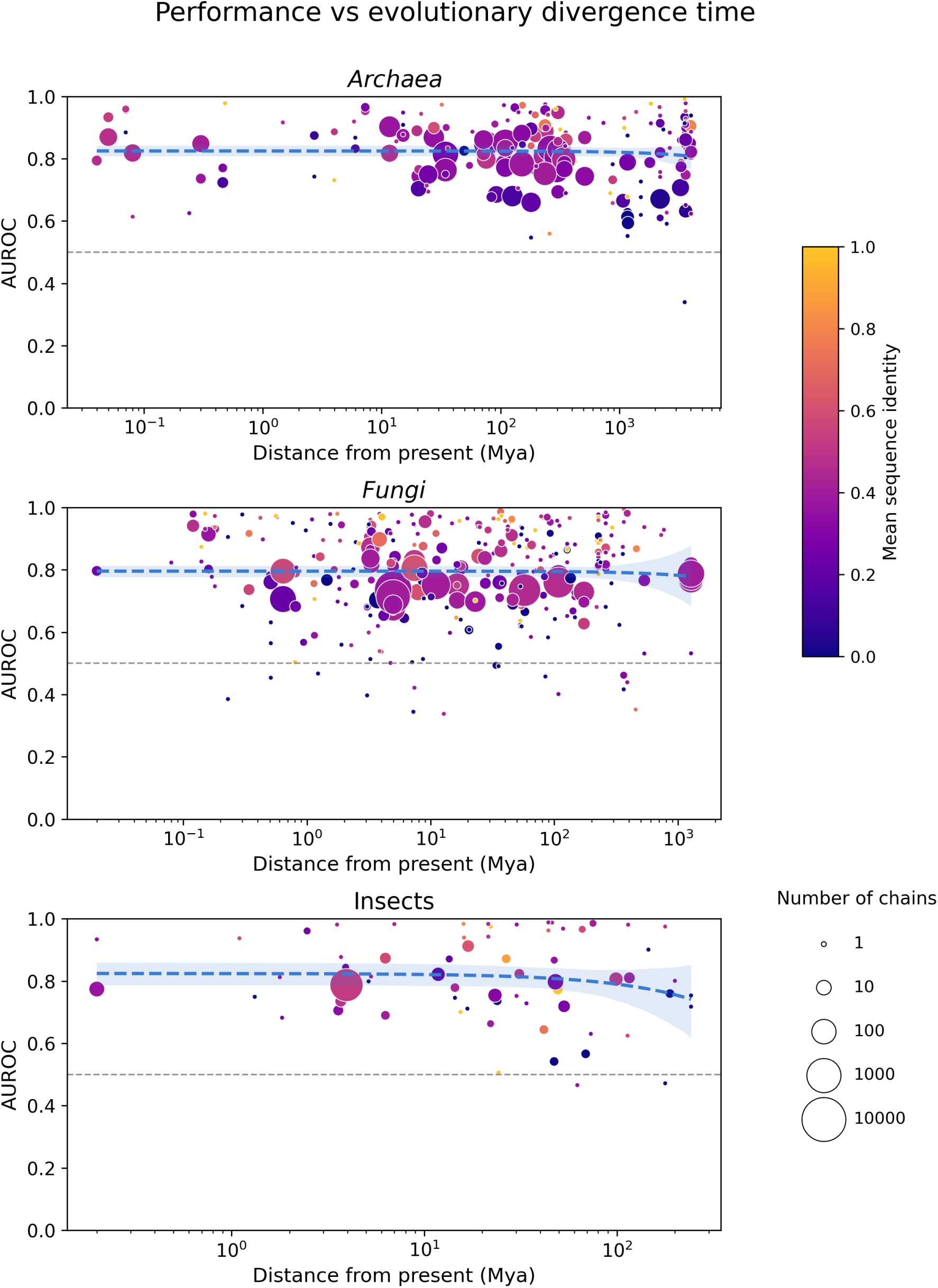
Species-level performance as a function of evolutionary divergence time for *Archaea*, *Fungi*, and *Insecta*. For each kingdom, the model was trained by excluding all protein pairs containing at least one protein from the corresponding taxonomic group, and subsequently evaluated on that held-out group. Each panel reports the mean AUROC per species as a function of evolutionary divergence time from the present (in millions of years). Points represent individual species; point size is proportional to the number of contributing chains, and colour encodes the mean sequence identity to the training set. Dashed lines indicate linear regression fits.

**Figure S7:**
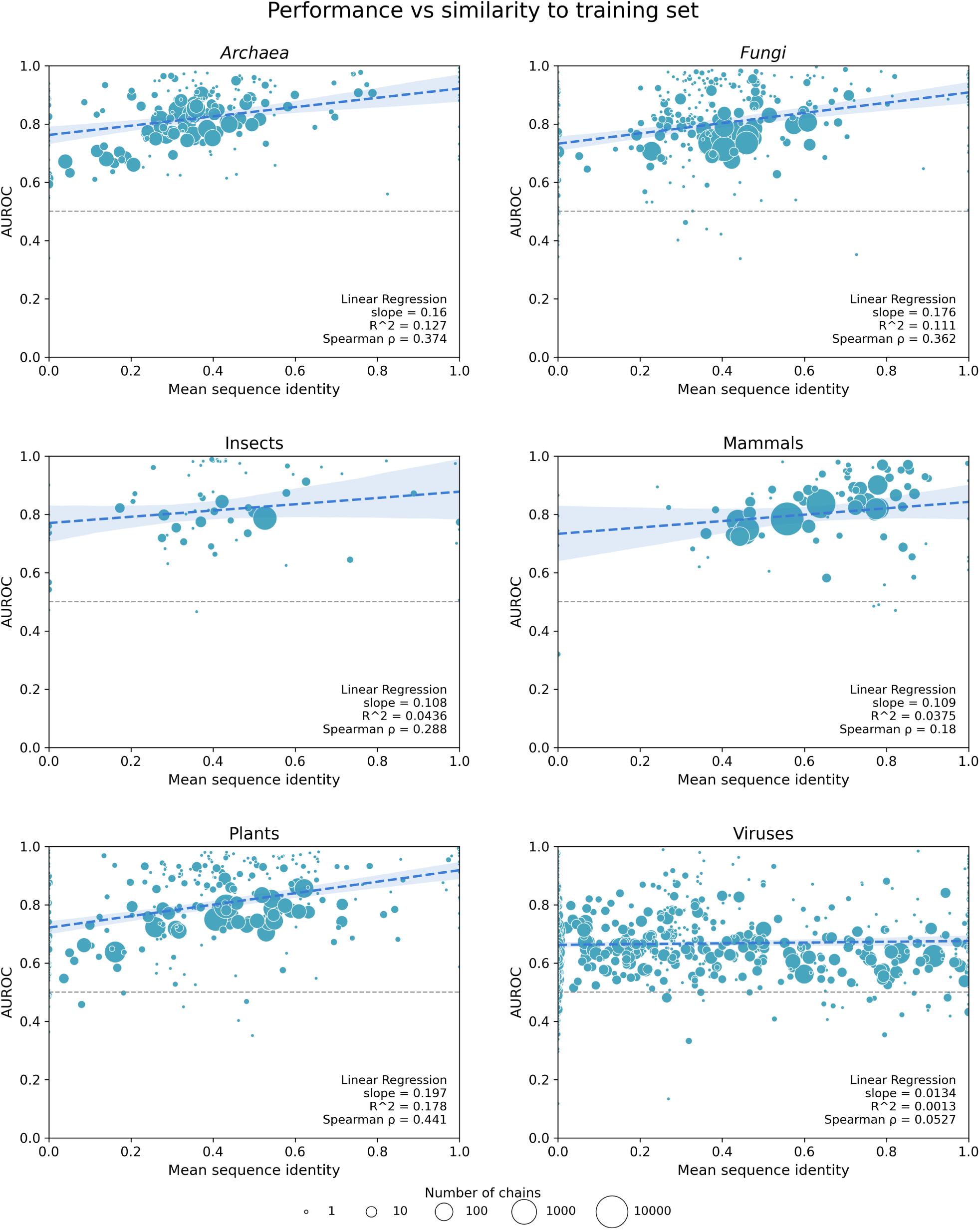
Species-level performance as a function of sequence identity for different taxonomic groups. (*Archaea*, *Fungi*, *Insecta*, *Mammalia*, *Viridiplantae*, and *Viruses*). For each group, the model was trained by excluding all protein pairs containing at least one protein from the corresponding taxonomic group, and subsequently evaluated on that held-out group. Each panel shows the mean AUROC per species as a function of the mean sequence identity to the training set. Points represent individual species, with size proportional to the number of contributing chains. Dashed lines indicate linear regression fits; the corresponding slope, *R*^2^, and Spearman’s rank correlation coefficient (*ρ*) are reported in each panel.

